# Integrin-deficient T cell leukemia accumulates in the central nervous system

**DOI:** 10.1101/2024.06.28.601274

**Authors:** Samantha Y. Lux, Cynthia Chen, Bibi S. Subhan, Hyunsoo Chung, Martyna Okuniewska, Asha Y. Caslin, Kathleen A. Martin, Jennifer K. Schiavo, Jonah B. Vernejoul, Robert C. Froemke, Michael Cammer, Susan R. Schwab

## Abstract

T-cell acute lymphoblastic leukemia (T-ALL) spreads aggressively to the central nervous system (CNS), particularly the leptomeninges. Children with T-ALL are treated with high-dose, CNS-directed chemotherapy, which can cause lasting neurotoxicity and is not always effective. Little is known about how T-ALL enters and persists within the CNS. However, normal T cell migration into the CNS has been extensively studied. Two integrins—VLA-4 and LFA-1—mediate normal T cell entry to the CNS, and VLA-4 blockade effectively treats multiple sclerosis by excluding T cells from the brain. We hypothesized that these integrins would likewise be required for T-ALL CNS entry. Unexpectedly, not only were VLA-4 and LFA-1 dispensable for T-ALL to reach the CNS, integrin-deficient T-ALL accumulated in the CNS compared to control. Mechanistically, integrin loss accelerated T-ALL proliferation in the CNS, suggesting that integrin-mediated interactions may promote quiescence in this space. Integrin blockade synergized with chemotherapy targeting proliferating cells, raising the possibility that combination therapy might be a powerful strategy.

## Introduction

Acute lymphoblastic leukemia (ALL) is the most common pediatric cancer [1, 2]. In recent years, major advances have improved the five-year survival rate to over 90% [2]. However, ALL’s propensity to infiltrate the central nervous system (CNS), particularly the meninges, remains a significant challenge [3, 4]. To eradicate disease, the standard-of-care is to treat all children diagnosed with ALL with high-intensity, CNS-directed chemotherapy [3, 5]. This regimen induces substantial toxicity and can cause irreparable neurological damage. Furthermore, despite aggressive treatment, the CNS is a frequent site of relapse. Our lack of understanding of how ALL enters and persists in the CNS has limited the development of targeted therapies for decades.

The T cell form of ALL, T-ALL, has especially high rates of CNS penetration [6]. Very little is known about how T-ALL infiltrates the CNS, but normal effector T cell trafficking into the CNS has been extensively characterized. T cells cross the blood-endothelial barrier into the CNS through a stepwise process of rolling, sticking, and transmigration [7]. This cascade requires integrins—heterodimeric adhesion molecules consisting of an alpha and beta chain—which allow T cells to attach firmly to the endothelium. The integrin VLA-4 (*α*4*β*1) is canonically required for T cells to enter the CNS [8, 9], and in some cases the integrin LFA-1 (*α*L*β*2) can contribute to this transmigration [7]. In fact, natalizumab, a monoclonal antibody blocking the alpha subunit of VLA-4, is widely used to treat multiple sclerosis by preventing autoreactive T cells from reaching the brain [10]. Additionally, integrins enable B-cell ALL to travel to the meninges via bridging channels between the skull bone marrow and the dura mater [11]. We hypothesized that blocking VLA-4 and LFA-1 would render T-ALL unable to metastasize to the CNS.

## Results

### T-ALL deficient in LFA-1 and VLA-4 preferentially accumulates in the CNS

To test whether T-ALL could enter the CNS in the absence of LFA-1 and VLA-4, we developed a T-ALL line by transducing the bone marrow from mice that lacked these integrins (double-knockout, DKO) with a constitutively active NOTCH-1, per an established protocol [12] (S. Fig. 1A). We chose a NOTCH1-driven model because mutations in NOTCH1 and its regulators are found in the majority of human T-ALL cases [13–15]. In the integrin-deficient T-ALL cells, the first and second exons of Itgal (αL, the alpha subunit of LFA-1) were knocked out, and the third exon of Itgb1 (β1, the beta subunit of VLA-4) was excised [16, 17]. Of note, B-cell ALL requires α6 integrin to enter the CNS, which we could not detect on the surface of DKO T-ALL (S. Fig. 1B,C). This is unsurprising as β1 integrin is a canonical partner of α6, and we could not detect mRNA or protein for the other canonical binding partner of α6 integrin, β4 integrin, in these cells (S. Fig. 1B,C; S. Table 1) [11, 18].

We co-transferred the DKO T-ALL cells at a 1:1 ratio with similarly generated NOTCH1-driven, mCherry^+^, wild-type (WT) T-ALL cells intravenously into sub-lethally irradiated WT adult mice (Fig. 1A,B). At a humane endpoint, we enumerated DKO and WT T-ALL cells in common sites of disease (Fig. 1C,D). As in human patients, CNS T-ALL cells in our model were largely confined to the leptomeninges rather than the brain parenchyma [4, 19]; washing the brain surface yielded 10-20 times more cells than extracting T-ALL cells from the brain parenchyma, and our quantification reflects leptomeningeal disease (S. Fig. 1D). Surprisingly, in the CNS we found nearly 10 times more integrin-deficient T-ALL cells than WT T-ALL cells. This was especially notable considering the DKO T-ALL’s relative absence in the spleen, blood and bone marrow, natural reservoirs for leukemia cells (Fig. 1C,D).

**Fig. 1.**
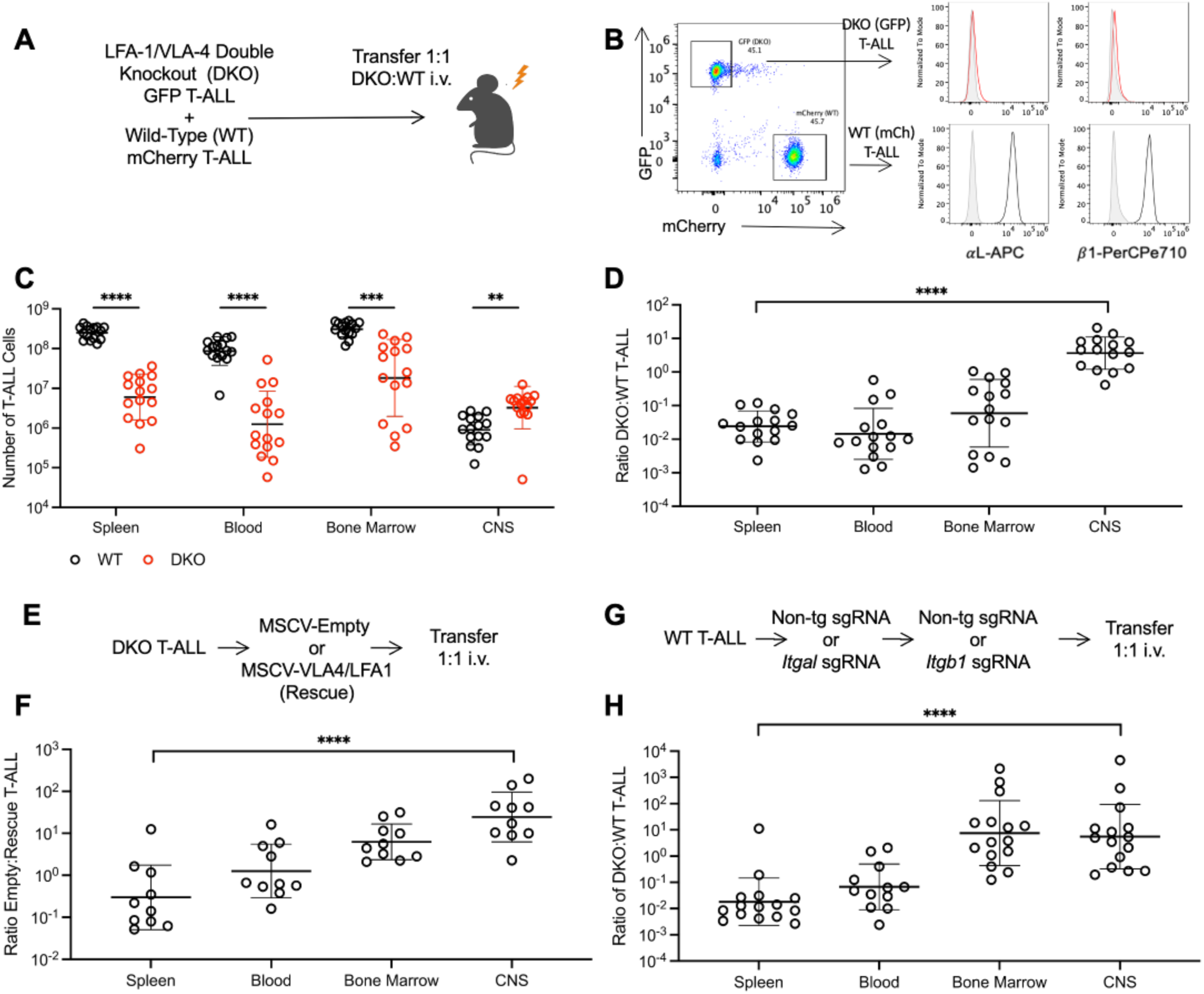
Integrin-deficient T-ALL accumulates in the CNS compared to WT T-ALL. **(A-D)** WT mCherry^+^ T-ALL and DKO GFP^+^ T-ALL cells were mixed 1:1 and transferred intravenously (i.v.) into sub-lethally irradiated recipients. **(A)** Experiment design. **(B)** Left: Representative flow cytometry plot distinguishing WT T-ALL and DKO T-ALL from the CNS. Right: Representative surface stain of β1 and αL integrins on T-ALL. Black: WT, red: DKO, grey: isotype. Numbers **(C)** and ratios **(D)** of DKO (red) and WT (black) T-ALL cells found in the indicated tissues at a humane endpoint. Data compiled from 3 experiments; *n* = 15. **(E-F)** DKO T-ALL cells were transduced with either vectors encoding *Itgal* and *Itgb1* (“rescue”) or empty vector (“empty”) and transferred 1:1 i.v. into sub-lethally irradiated recipients. **(E)** Experiment design. **(F)** Ratio of DKO-empty to DKO-rescue T-ALL cells in the indicated tissues at a humane endpoint. Data compiled from two experiments; *n* = 10. **(G-H)** WT T-ALL cells were nucleofected with sgRNAs targeting *Itgal* or non-targeting gRNAs along with recombinant Cas9 protein. The process was repeated for *Itgb1* to generate *“*CRISPR-DKO” and “CRISPR-WT T-ALL.” The cells were mixed 1:1 and transferred i.v. into sub-lethally irradiated recipients. **(G)** Experiment design. **(H)** Ratio of CRISPR-DKO to CRISPR-WT T-ALL cells found in the indicated tissues at a humane endpoint. Data compiled from five experiments; *n* = 15.

To determine whether DKO T-ALL accumulation in the CNS was solely due to the absence of LFA-1 and VLA-4, we tested whether restoration of VLA-4 and LFA-1 in DKO T-ALL would reverse its accumulation in the CNS. We transduced the DKO T-ALL line with vectors encoding *Itgal* and *Itgb1* or an empty vector to generate “rescue” and “empty” lines (Fig. 1E). We found that restoration of αL and β1 in the DKO T-ALL (DKO-rescue) reduced the number of cells accumulating in the CNS compared to vector-transduced cells (DKO-empty) (Fig. 1F).

To further test whether DKO T-ALL accumulation in the CNS was solely due to the absence of LFA-1 and VLA-4, we used CRISPR-Cas9 to target the first through third exons of *Itgal* and the first and second exons of *Itgb1* in the WT T-ALL line (Fig. 1G, S. Fig. 1E,F). These CRISPR-DKO T-ALL lines recapitulated the original phenotype, with CRISPR-DKO T-ALL preferentially accumulating in the CNS compared to control-targeted cells (CRISPR-WT) (Fig. 1H).

We saw similar results when the DKO and WT T-ALL were transferred singly (S. Fig. 2A, B). Additionally, irradiation of the CNS was not required for DKO T-ALL to accumulate in the CNS (S. Fig. 2C). Taken together, these data suggest that the loss of VLA-4 and LFA-1 results in T-ALL accumulation within the CNS.

### Integrin loss is unlikely to induce T-ALL accumulation in the CNS by enhancing CNS entry, limiting CNS exit, or promoting immune escape

Having disproven our original, seemingly obvious, hypothesis, we sought to explain how loss of integrins resulted in T-ALL accumulation in the CNS. It was possible that loss of integrins paradoxically improved entry into the CNS, limited exit from the CNS, or promoted T-ALL survival or proliferation within the CNS.

#### Entry: Premature release from bone marrow

One explanation for our results was that loss of integrins led to premature release of T-ALL cells from the bone marrow, a key T-ALL niche. Integrins retain normal hematopoietic progenitor cells in the bone marrow; when VLA-4 binding is blocked, these cells exit their niche and move into the blood [20–22]. We hypothesized that this may be occurring in T-ALL, thus allowing DKO to migrate earlier to the CNS and take over the leptomeningeal space. To test this, we conducted a time-course experiment, assessing the ratio of DKO:WT T-ALL in the CNS at different stages of disease. Contrary to our expectation, we found that WT T-ALL accumulated in the CNS before DKO T-ALL. At around mid-stage disease, DKO T-ALL caught up, and at late-stage disease, DKO T-ALL overtook WT leukemia (Fig. 2A, B).

**Fig. 2.**
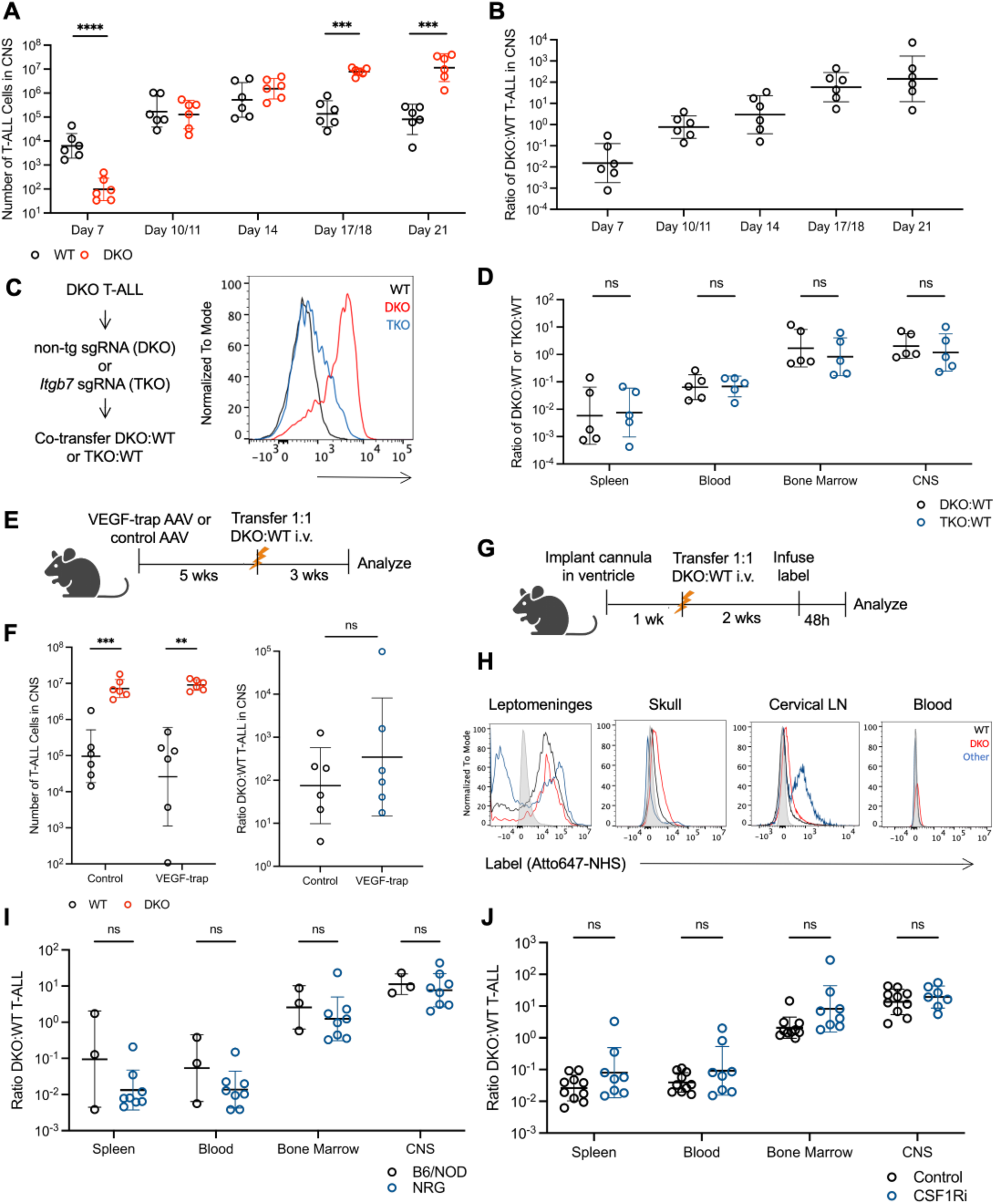
Integrin-deficient T-ALL is unlikely to accumulate in the CNS due to enhanced entry, limited exit, or superior immune escape. **(A, B)** Sub-lethally irradiated WT mice were injected i.v. with 1:1 DKO:WT T-ALL and tissues were isolated from 3 mice every 3-4 days. **(A)** Total number of WT (black) and DKO (red) T-ALL cells in the CNS. **(B)** Ratio of DKO:WT T-ALL in the CNS. Data compiled from two experiments; *n* = 36. **(C)** Left, CRISPR-TKO T-ALL generation. Right, representative surface staining of integrin *β*7 on T-ALL cells (blue: TKO, black: WT, red: DKO). **(D)** Ratio of DKO:WT T-ALL (black) or TKO:WT T-ALL (blue) in indicated tissues at a humane endpoint. Data compiled from two experiments; n = 5. **(E, F)** WT mice were infected with AAV encoding a VEGF-trap or control AAV. 5 weeks later, mice were sub-lethally irradiated and injected i.v. with 1:1 DKO:WT T-ALL. At a humane endpoint, tissues were isolated and T-ALL was enumerated. **(E)** Experiment design. **(F)** Number of WT (black) and DKO (red) T-ALL or ratio of DKO:WT T-ALL in the CNS of mice treated with either control AAV or AAV-VEGF-trap. Data compiled from 2 experiments; n = 6. **(G, H)** WT mice were surgically implanted with an intra-cerebroventricular cannula. 1 week later, mice were sub-lethally irradiated and injected i.v. with 1:1 DKO:WT T-ALL. At mid-stage disease, a reactive label (Atto647-NHS-ester) was infused through the cannula into the left lateral ventricle. 48h later, tissues were isolated and labelled T-ALL cells were enumerated. **(G)** Experiment design. **(H)** Representative flow cytometry plots of Atto647 labelling in the indicated tissues. WT T-ALL, black; DKO T-ALL, red; other leukocytes, blue. Representative of 3 experiments; *n* = 6. **(I)** NRG mice or WT controls (B6 or NOD) were sub-lethally irradiated, injected i.v. with 1:1 DKO:WT T-ALL, and analyzed at a humane endpoint. Ratio of DKO:WT T-ALL cells in controls (black; *n* = 3) or NRG mice (blue; *n* = 8). Data compiled from 2 experiments. **(J)** WT mice were fed either diet supplemented with a CSF1R inhibitor (PLX5622) or control diet for 1 week, sub-lethally irradiated and injected i.v. with 1:1 DKO:WT T-ALL, and maintained on either PLX5622 or control diet until a humane endpoint. Ratio of DKO:WT T-ALL in the indicated tissues of mice fed control diet (black; *n* = 10) or PLX5622 diet (blue; *n* = 8). Data compiled from 2 experiments. Bars represent geometric mean and geometric SD. Statistics calculated on log-transformed data using unpaired t-test with Welch’s correction. *p ≤ 0.05, **p ≤ 0.01, ***p ≤ 0.001, ****p ≤ 0.0001

#### Entry: Up-regulation of other integrins

A second attractive explanation for our results was that loss of VLA-4 and LFA-1 resulted in compensatory expression of other integrins that were better matched to adhesion molecules expressed by CNS blood endothelial cells in the context of leukemia. In fact, when we performed bulk RNA sequencing comparing DKO and WT T-ALL isolated from the CNS, we found several differences in integrin expression between DKO and WT T-ALL (S. Fig. 2D,E). Of these, only *Itgb7* was significantly increased in DKO compared to WT T-ALL. We confirmed elevated surface protein expression of β7 integrin (Fig. 2C), which might reflect a combination of increased mRNA and increased availability of its binding partner *α*4 integrin upon the loss of β1 integrin. The ligand for α4β7 integrin, MAdCAM1, has primarily been characterized for its role in T cell migration into the gut [23], but has also been found to be expressed in the choroid plexus during inflammation [24]. Furthermore, α4β7 integrin has been reported to facilitate T cell entry into the CNS in some settings [25].

To test whether DKO T-ALL was using α4β7 to cross into the CNS, we triply knocked out αL, β1, and β7 (third exon) in T-ALL cells using CRISPR-Cas9 as described above (Fig. 2C). We found that triple-knockout (TKO) T-ALL was indistinguishable from DKO T-ALL in its accumulation in the CNS (Fig. 2D). These results suggest that integrin α4β7 is not responsible for the increased accumulation of DKO T-ALL within the CNS.

#### Exit

We next hypothesized that DKO T-ALL may accumulate in the CNS due to a defective ability to exit the CNS. The dorsal lymphatics of the meningeal dura mater have been recently characterized as a key route of exit for lymphocytes from the CNS to the draining cervical lymph nodes [26, 27], and integrins in some cases enable T cells to access lymphatics [28, 29].

If DKO T-ALL accumulated in the CNS due to a failure to exit through the dorsal meningeal lymphatics, the ratio of DKO:WT T-ALL cells in the CNS should normalize in the absence of meningeal lymphatics. To ablate the meningeal lymphatics, we took advantage of the fact that, unlike most lymphatic vessels in the body, the dorsal meningeal lymphatics require continuous VEGFR3 signaling to persist [30]. We therefore treated mice with a “VEGF-trap,” an adeno-associated virus encoding the ligand-binding domain of VEGFR3 fused to IgG-Fc, which captures available VEGFC/D and induces dorsal meningeal lymphatic regression. As a control, we treated mice with an adeno-associated virus that encodes the stalk (non-ligand-binding) of VEGFR3 fused to IgG-Fc [30]. Five weeks later, we induced leukemia in these animals by co-transferring WT and DKO T-ALL (Fig. 2E). We found that the regression of meningeal lymphatics had no effect on the ratio of the DKO to WT T-ALL in the CNS, or on the overall number of cells in the CNS (Fig. 2F). Meningeal lymphatic regression was confirmed by microscopy (S. Fig. 3A). We found similar results when we repeated this experiment using mice that were developmentally lacking in meningeal lymphatics (mice expressing the VEGF-trap under the keratin-14 promoter) [27] (S. Fig. 3B,C). These data suggest that DKO T-ALL does not preferentially accumulate in the CNS due to a requirement for integrins to exit through the meningeal lymphatics.

There may be alternate, uncharacterized pathways of exit from the CNS that could require integrins. To address more generally whether DKO T-ALL cells were defective in CNS egress compared to WT T-ALL cells, we labeled T-ALL cells in the CNS and assessed their location 48 hours later. Mice were surgically implanted with an intraventricular cannula in the lateral ventricle of the CNS prior to leukemia induction. Once mice reached mid-stage disease, we infused a reactive, fluorescent dye into the CNS via the cannula (Fig. 2G). 48 hours later, we found that T-ALL cells in the CNS were efficiently labeled. We found both labeled and non-labeled normal leukocytes in the CNS and the draining cervical lymph nodes, suggesting that normal leukocytes were cycling through the CNS. However, we did not detect labeled T-ALL cells in any other site, including blood, skull bone marrow, or the CNS-draining cervical lymph nodes (Fig. 2H). Overall, we found no evidence that delayed exit causes accumulation of DKO T-ALL within the CNS.

#### Survival: Immune escape

As differences in trafficking seemed unlikely to explain the CNS accumulation of the DKO T-ALL, we next hypothesized that differential interactions with the host immune system might be responsible. Integrins play a key role in cementing synapses between leukocytes [31], and cytotoxic T cells can have a powerful anti-tumor effect in ALL [32, 33]. Perhaps the lack of integrins enabled the DKO T-ALL to escape immune surveillance.

If reduced susceptibility to an anti-leukemia immune response explained DKO T-ALL accumulation within the CNS, the ratio of DKO to WT T-ALL in the CNS should normalize in an immune-deficient host. To test this, we co-transferred WT and DKO T-ALL cells into NOD-*Rag1^-/-^Il2rg^-/-^*(NRG) mice, which lack B, T, and NK cells, but found no difference in the ratio of the DKO:WT T-ALL in the CNS between the NRG mice and either WT C57BL/6 or NOD controls (Fig. 2I). Interestingly, the B6-derived leukemia was frequently cleared entirely in immune-competent NOD hosts (not shown). T-ALL cells also interact with myeloid cells, although integrin-mediated interactions with myeloid cells promote T-ALL progression in many tissues [34, 35]. We tested a role of myeloid cells in the CNS accumulation of DKO cells by treating mice with the colony-stimulating factor 1 receptor (CSF1R) inhibitor PLX5622, which has been widely used to deplete microglia [36] as well as meningeal macrophages [37, 38], prior to induction of leukemia. We again found no difference in the ratio of DKO:WT T-ALL in the CNS between mice treated with the CSF1R inhibitor and controls (Fig. 2J). Consistent with previous findings that myeloid cells support T-ALL persistence in the CNS, we found that treatment with the CSF1R inhibitor resulted in an overall reduction in the number of T-ALL cells (S. Fig. 3D-F) [39]. More generally, we did not observe a difference in the rate of apoptosis between DKO and WT T-ALL in the CNS, as assessed by flow cytometry of cells stained for cleaved caspase 3 (S. Fig. 3G,H). Taken together, these data suggest that DKO T-ALL is not accumulating in the CNS because of its avoidance of an anti-leukemia immune response.

### The loss of integrins in T-ALL confers a proliferation advantage in the CNS

#### DKO T-ALL proliferates more rapidly than control T-ALL in the CNS

Within the dura mater, we observed that the DKO and WT T-ALL localized differently—the DKO cells tended to spread through the tissue while the WT cells tended to cluster near blood vessels (Fig. 3A). This is consistent with observations of increased effector T cell motility within the spinal cord meninges upon intrathecal injection of integrin-blocking antibodies [40].

**Fig. 3:**
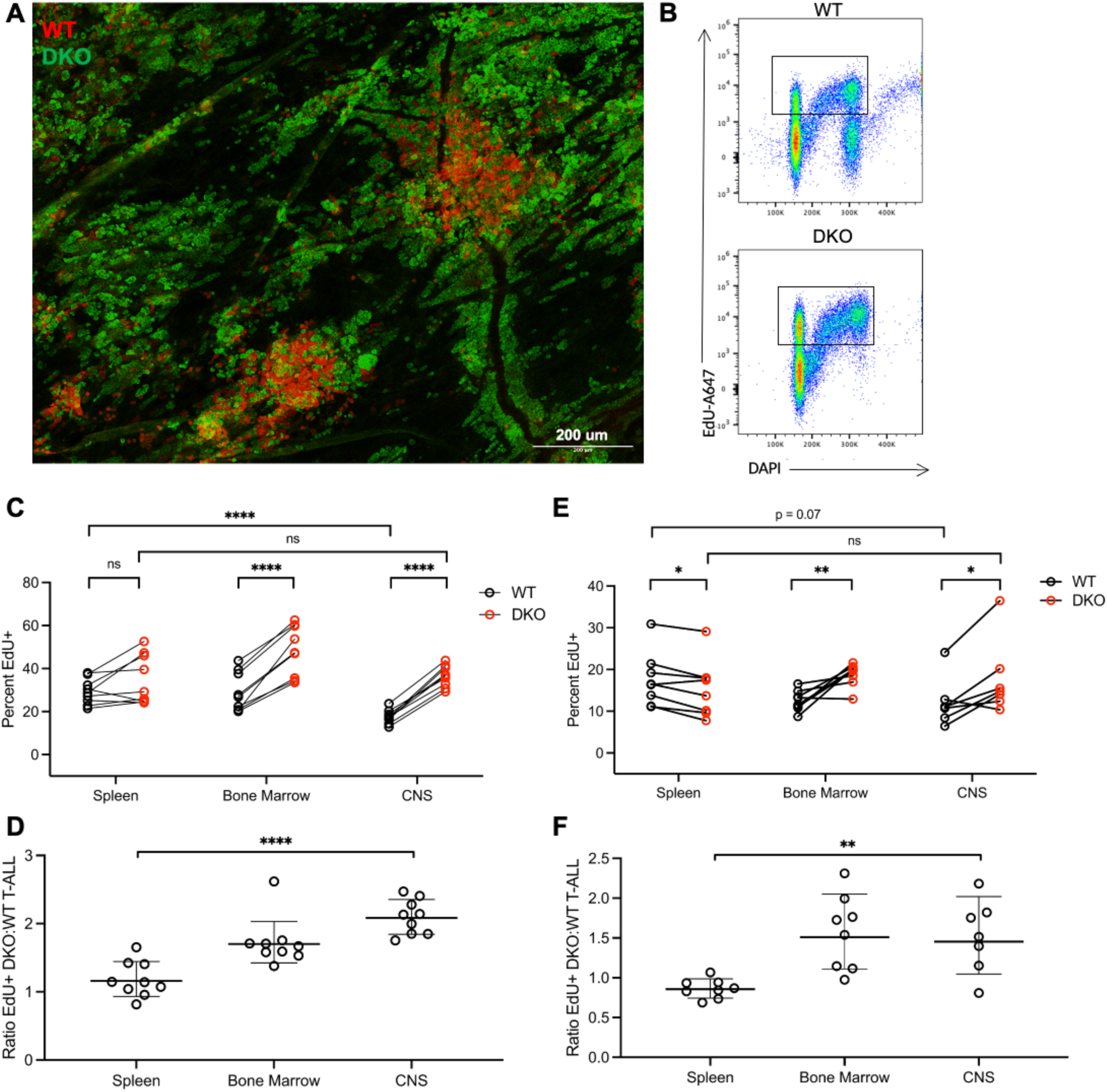
Integrin-deficient T-ALL proliferates more than WT T-ALL in the CNS. **(A)** Representative (of *n* = 12) immunofluorescence image of whole-mounted dural meninges of mice injected i.v. with 1:1 DKO:WT T-ALL. Mice were analyzed at mid-stage disease. Red, WT T-ALL; green, DKO T-ALL; scale bar, 200*µ*m. **(B)** Representative flow cytometry gating to identify proliferating cells isolated from tissues of mice that received EdU 4 hours before analysis (in two intraperitoneal doses, 100 *µ*g at -4h and 100 *µ*g at -2h). **(C, D)** Sub-lethally irradiated WT mice received 1:1 DKO:WT T-ALL cells i.v., and were analyzed at mid-stage disease. **(C)** The percent EdU+ of WT (black) or of DKO (red) T-ALL cells and **(D)** the ratio of % EdU+ DKO: % EdU+ WT T-ALL cells in the indicated tissues. Data compiled from 3 experiments; *n* = 9. **(E, F)** As in (C, D) but for the CRISPR-WT and CRISPR-DKO pair of T-ALL lines. Data compiled from 4 experiments; *n* = 8. For (C, E), statistics comparing WT and DKO within a tissue were calculated on logit-transformed data using paired t-test. Statistics comparing WT or DKO from CNS to spleen were calculated on logit-transformed data using unpaired t-test with Welch’s correction. For (D, F), statistics calculated using unpaired t-test with Welch’s correction. Bars represent geometric mean and geometric SD. *p ≤ 0.05, **p ≤ 0.01, ***p ≤ 0.001, ****p ≤ 0.0001

We hypothesized that a difference in distribution might result in differential access to nutrients or differential signals from the stromal niche and in turn affect proliferation. Indeed, interactions with meningeal stroma have been shown to limit ALL proliferation in culture and xenotransplants [41]. We therefore performed an *in vivo* pulse-labelling experiment with 5-ethynyl-2′-deoxyuridine (EdU) and assessed the incorporation of EdU and the abundance of DNA (cell cycle stage) by flow cytometry (Fig. 3B). We observed proliferating DKO and WT T-ALL cells in the CNS, bone marrow, and spleen, and found that DKO T-ALL was more proliferative than WT in both the bone marrow and CNS (Fig. 3C). Furthermore, the greatest difference between DKO and WT T-ALL proliferation was present in the CNS (Fig. 3D). When we repeated this experiment with the CRISPR-DKO and WT T-ALL pair, we again observed increased proliferation of DKO T-ALL in the CNS (Fig. 3E,F). Differences between the DKO and WT T-ALL in other tissues were less consistent between the two pairs of lines. Interestingly, when comparing the proliferation rate of either DKO or WT T-ALL across tissues, there was little difference in the DKO T-ALL proliferation rate between spleen, bone marrow, and CNS; rather, the proliferation of the WT T-ALL was suppressed in the CNS when compared to spleen (Fig. 3C,E).

We hypothesized that the unique stroma of the CNS might be responsible for suppressing the proliferation of WT T-ALL. Fibroblasts are abundant in the meninges and can powerfully modulate immune responses [42–47]. We expanded dural fibroblasts from mice with T-ALL and assessed their transcriptome by RNA-sequencing [43]. We used the NicheNet program to match ligands expressed by the fibroblasts with altered signaling in T-ALL indicated by differentially expressed genes in control versus DKO T-ALL isolated from the CNS [48, 49]. NicheNet highlighted TGF-beta as most likely to be responsible for differences between control and DKO T-ALL, followed by oncostatin M (S. Fig. 4A, B). Future work will test these candidates.

### Integrin blockade synergizes with chemotherapy targeting proliferating cells to limit CNS T-ALL

One of the mainstays of therapy for ALL is to target rapidly-proliferating cells with nucleotide-synthesis inhibitors [3]. We hypothesized that if increased proliferation of DKO compared to WT T-ALL cells in the CNS was in fact responsible for the increased accumulation of DKO compared to WT T-ALL in the CNS, the ratio of DKO:WT T-ALL should normalize if proliferation was limited. To test this, we treated mice at mid-stage disease with the thymidylate synthase inhibitor 5-flurouracil (5FU), which reaches the cerebrospinal fluid, for three days [50–53] (Fig. 4A). We found that DKO T-ALL was depleted more than WT T-ALL in the CNS, where the ratio of DKO:WT T-ALL became nearly 1:1 in just three days (Fig. 4B). When we repeated this experiment with the CRISPR-DKO and WT T-ALL pair, we again observed a reduction in the ratio of DKO:WT T-ALL in the CNS with a short treatment (Fig. 4C). The efficacy of 5FU treatment was confirmed by enumerating total T-ALL in key tissues compared to vehicle-treated mice (S. Fig. 4C,D).

**Fig. 4:**
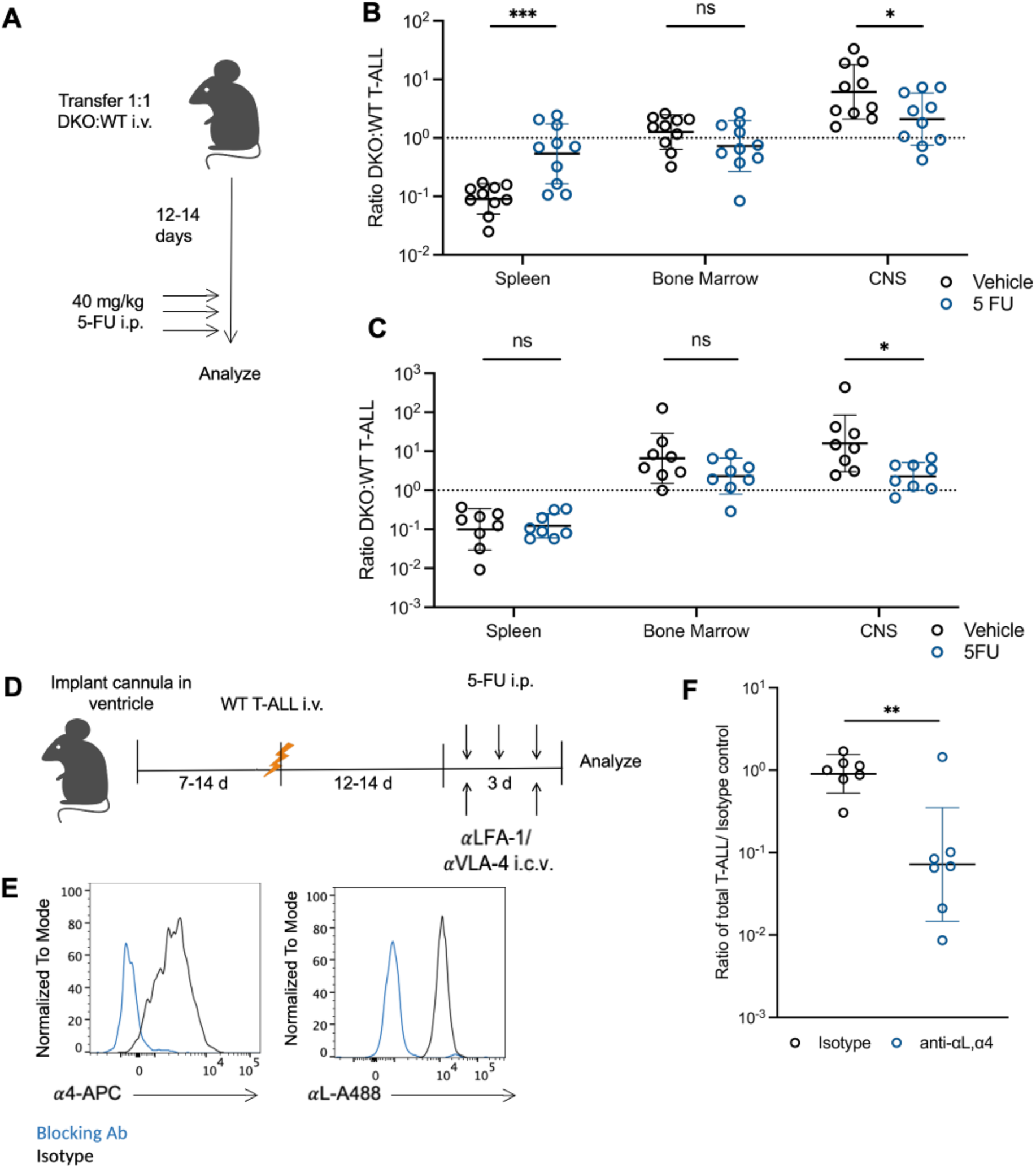
Integrin-deficient T-ALL is more susceptible to chemotherapy targeting dividing cells and integrin blockade synergizes with chemotherapy to limit CNS T-ALL. **(A-C)** Sub-lethally irradiated WT mice received 1:1 DKO:WT T-ALL i.v., were treated daily i.p. with 40mg/kg 5-fluorouracil (5-FU) for 3 days at mid-stage disease (starting 12-14 days after T-ALL transfer), and were analyzed 24h after the last dose of 5-FU. **(A)** Experiment design. **(B)** The ratio of DKO:WT T-ALL cells in vehicle-treated (black; *n* = 10) or 5FU-treated (blue; *n* = 10) mice. Data compiled from 2 experiments. **(C)** As in (B) but for the CRISPR-WT and CRISPR-DKO pair of T-ALL lines. Vehicle-treated (black; *n* = 8), 5FU-treated (blue; *n* = 8). Data compiled from 2 experiments. **(D-F)** Mice were surgically implanted with a cannula in the left lateral ventricle. At least 7 days later, they were sub-lethally irradiated and injected i.v. with WT T-ALL. At mid-stage disease (12-14 days after T-ALL transfer), mice were injected i.p. with 40mg/kg 5-FU daily for three days. On days 1 and 3 of 5-FU treatment, mice received 5 microliter intraventricular infusions of isotype (47 microgram) or anti-LFA-1 and anti-VLA-4 (26 micrograms each) blocking antibodies. 24h after the last dose, tissues were isolated and T-ALL cells were enumerated. **(D)** Experiment design. **(E)** Surface expression of *α*4 and *α*L integrins on WT T-ALL from the CNS after intra-cerebroventricular injection of LFA-1- and VLA-4- blocking antibodies (blue; representative of *n* = 7) or isotype (black; representative of *n* = 7). Representative of 4 experiments. **(F)** Total number of WT T-ALL in the CNS normalized to the average isotype control per experiment. Data compiled from 4 experiments; *n* = 7 pairs. Bars represent geometric mean and geometric SD. Statistics calculated on log-transformed data using unpaired t-test with Welch’s correction. *p ≤ 0.05, **p ≤ 0.01, ***p ≤ 0.001, ****p ≤ 0.0001

Given these findings, we hypothesized that in wild-type T-ALL disease, the efficacy of chemotherapy in the CNS could be improved by simultaneously blocking integrins; this strategy might encourage T-ALL cells to proliferate and thereby render them sensitive to the drugs. We used an intraventricular cannula to administer integrin-blocking antibodies to the CNS of mice with established WT T-ALL at the same time that they received systemic chemotherapy (5FU) (Fig. 4D). We confirmed the efficacy of blocking antibodies by flow cytometry (Fig. 4E). We found that the combination of intracranial blocking antibodies and systemic chemotherapy was far more effective at reducing total CNS T-ALL than chemotherapy alone (Fig. 4F). Mice that received intracranial antibodies without chemotherapy had no significant difference in CNS T-ALL accumulation from those that received isotype (S. Fig. 4E); however, administration of antibodies without reducing the T-ALL burden with 5FU resulted in a high rate of mortality (3/8 vs 1/16).

## Discussion

Much to our surprise, the loss of LFA-1 and VLA-4 in T-ALL did not block leukemia access to the CNS but instead promoted T-ALL accumulation in the CNS. It now seems improbable that T-ALL enters the CNS by crossing an intact endothelial barrier. Perhaps T-ALL either damages the endothelium to enter the CNS or migrates into the CNS via adjacent tissues, such as the bridging channels linking the skull bone marrow to the dura mater, although B-ALL transit through the bridging channels has been shown to require integrins that T-ALL does not express in our model [11]. An important future question is how T-ALL accesses the CNS: what route does it take into the meninges, and what are the mechanical requirements for this migration? Mechanistically, integrin expression suppresses proliferation of T-ALL within the CNS, likely by mediating interactions with the meningeal stroma. A second key question is how integrins limit T-ALL proliferation in the CNS. The CNS is a canonically immune-suppressive environment, and the nature of CNS immune responses is an area of active research[42, 47, 54, 55]. We hypothesize that interactions between the meningeal stroma and normal T cells may limit over-exuberant responses and incidentally restrain T-ALL division.

Finally, these findings have important implications for the design of therapeutic strategies for ALL. We have found strong synergy between integrin blockade and anti-proliferative chemotherapy to treat CNS T-ALL. An analysis of 261 blood and bone marrow biopsies from pediatric and young adult T-ALL patients revealed consistently high integrin expression [13] (S. Fig. 5D). High integrin expression by B-ALL cells independently predicts poor prognosis [35, 56–59], and VLA-4 expression has been found to mediate chemoresistance of ALL in the bone marrow [58, 60–64]. Within the CNS, proteomic and transcriptomic analyses of ALL indicate altered expression of adhesion proteins [65–67]. Furthermore, upon xenotransplantation of Jurkat T-ALL cells into mice, combined treatment with the chemotherapeutic drug cytarabine and Tris[2-(dimethylamino)ethyl]amine, which disrupts adhesion of Jurkat cells to meningeal cells in culture, reduced the number of Jurkat cells in the meninges compared to cytarabine alone [41]. Our findings highlight the need to understand what the fate of leukemia cells is when they are untethered from their niche. This might vary drastically from tissue to tissue, and understanding these cell dynamics will inform the optimal combination of treatments. Thus, for example, integrin loss alone was toxic in our model, but integrin blockade was highly effective in combination with 5FU. Ultimately, targeting integrins in T-ALL may be a powerful strategy to reduce the need for prolonged neurotoxic therapies.

## Materials and methods

### Animals

C57BL/6 (CD45.2), B6.SJL-PtprcaPepcb/BoyJ (CD45.1), Mx1-cre [68], *Itgb1*^f/f^ [17], *Itgal*^-/-^ [16] and NRG (NOD background, *Rag1*^-/-^*Il2rg*^-/-^ [69]) mice were obtained from Jackson Laboratory. K14-VEGFR3-Ig mice were kindly provided by Kari Alitalo [70]. Mice were compared to littermate controls, unless otherwise specified. For CD45.2 or CD45.1/.2 hosts, female adult mice (6-12 weeks) were used. For all other hosts, male and female adult mice (6-12 weeks) were used depending on availability, as sex did not seem to affect the results. Mice were housed in specific pathogen-free conditions at New York University School of Medicine animal facilities. All cages were on a 12-h light/dark cycle (light 06:00-18:00) in a temperature-controlled and humidity-controlled room. Room temperature was maintained at 72 ± 2 °F (22.2 ± 1.1 °C), and room humidity was maintained at 30–70%. All animal experiments were performed in accordance with protocols approved by the New York University Grossman School of Medicine Institutional Animal Care and Use Committee.

### Bone marrow transduction and transplantation for generation of T-ALL

To generate the original GFP+ DKO and mCherry+ WT T-ALL lines, hematopoietic cells were isolated from bone marrow by flushing femurs and tibias. Stem and progenitor cells were enriched by magnetic selection of cells expressing c-Kit (STEMCELL EasySep Mouse Biotin Positive Selection Kit II). Cells were cultured in the presence of 50 ng/mL Flt3 ligand, 50 ng/mL stem cell factor (SCF), 10 ng/mL interleukin-3 (IL-3), and 10 ng/mL interleukin-6 (IL-6) (PeproTech) in OptiMEM + GlutaMAX (Invitrogen) supplemented with 10% fetal bovine serum (FBS)(Cytiva), 100 IU/mL penicillin and 100 ug/mL streptomycin (Gibco or Corning), and 55 μM β-mercaptoethanol (Sigma). 24 and 48 hr after enrichment, c-Kit+ cells were infected with retrovirus (MSCV Notch1ΔE-IRES-GFP or Notch1ΔE-IRES-mCherry), generated by transfection of Platinum E (Plat-E) retroviral packaging cells (Cell BioLabs Inc.), and concentrated by filtration according to the manufacturer’s instructions (Amicon Ultra-15; 100,000 NMWL). Transduction efficiency was determined by reporter fluorescence at 96 hr. Irradiated C57BL/6 mice (two rounds of 500 rads) received 35 * 10^3^ Notch1-ΔE-IRES-GFP/mCherry+, lineage− (lineage markers: CD4, CD8, B220, Gr1, Ter119, Cd11b) cells, together with 500 * 10^3^ unfractionated bone marrow cells for hemogenic support, intravenously by retro-orbital injection. Mice were bled regularly to monitor leukemia progression. The DKO donor hematopoietic cells were *Itgal*^-/-^ *Itgb1*^f/f^ *Mx1*-Cre. To delete *Itgb1*, when GFP+ blasts reached >10% of peripheral blood, mice received three intraperitoneal injections of poly(I:C) (10 ug/g, GE Heathcare) in PBS, administered every other day. Notch1ΔE-IRES-GFP *Itgb1^f/f-^ Itgal^-/-^ Mx1-cre* cells (DKO T-ALL) or Notch1ΔE-IRES-mCherry cells (WT T-ALL) were isolated from bone marrow at a humane endpoint.

*For secondary model induction* T-ALL cells were maintained *in vitro* (see below). For induction, recipients were sub-lethally irradiated (550 cGy) and retro-orbitally injected with 1–2 * 10^6^ T-ALL cells. Radiation was performed with either Cs or X-ray irradiator (X-rad 320/350).

### T-ALL cell culture

Primary mouse T-ALL cells isolated from bone marrow were maintained on OP9 stromal cells [71], kindly provided by Iannis Aifantis, in OptiMEM + GlutaMAX (Invitrogen) supplemented with 10% fetal bovine serum (FBS), 5 ng/ml IL-7, 100 IU/mL penicillin and 100 ug/mL streptomycin (Gibco or Corning), and 55 μM β-mercaptoethanol. Cells were passaged every 3–4 days onto a fresh feeder layer. OP9 cells were cultured in 75cm^2^ flasks in MEMα + GlutaMAX supplemented with 20% FBS and 100 IU/mL penicillin and 100 ug/mL streptomycin (Gibco or Corning), and passaged every 2-3 days. For co-culture, OP9s were pre-treated with 7.5 ng/mL mitomycin C (Sigma) for 3-4 hours to prevent feeder cell division. T-ALL cells for leukemia induction were isolated from co-culture by density gradient (Ficoll-paque Premium, Cytiva).

### Retroviral transduction of T-ALL cells

The coding regions of *Itgb1* and *Itgal* (Dharmacon) were cloned into the pMXs-IRES-mCherry retroviral vector, and Plat-E packaging cells were used for the generation of retrovirus. DKO T-ALL cells, seeded on mitomycin C-treated OP9s, were transduced with a retrovirus encoding either *Itgb1* or *Itgal* or empty vector. 48-72 hours after transduction, cells were sorted on a BD FACSAria II for positive surface expression of αL or β1. Cells were allowed to recover on MMC-treated OP9s for 48 hours and another round of transduction was performed using the plasmid carrying the other gene. After 48-72 hours, T-ALL cells double-positive for surface expression of αL and β1 were sorted on the BD FACSAria II.

### CRISPR deletion of genes in T-ALL cells

gRNAs targeting *Itgal*, *Itgb1* and *Itgb7* were ordered from IDT (sequences below) for use with the IDT Alt-R CRISPR-Cas9 system. ALT-R CRISPR-Cas9 Negative Control crRNA#1 was ordered as a non-targeting control (IDT). On day of nucleofection, gRNAs were resuspended in sterile tris-EDTA pH 8.0 to a concentration of 100uM. 1.2 uL of each gRNA for either *Itgal* or *Itgb1* or *Itgb7* were mixed with 1.7 uL of Alt-R S.p. HiFi Cas9 Nuclease V3, and 0.9 uL of PBS (total volume of 5 uL), and incubated for 20 minutes at room temperature, per manufacturer’s instructions. 1-2*10^6^ T-ALL cells were resuspended in 100uL P3 nucleofector solution (82 uL nucleofection solution + 18 uL supplement; Lonza P3 Primary Cell 4D-Nucleofector^TM^ X Kit), combined with gRNA-Cas9 complex, and nucleofected using program “*activated T cells”* using the Lonza Amaxa 4D Nucleofector. Immediately after, 500 uL warm media was added to cells in cuvette and incubated for 10 min at 37C before cells were returned to culture. After 48-72 hours, T-ALL cells were sorted based on surface expression of αL or β1 on BD FACSAria II. T-ALL cells were then subjected to a second round to delete both *Itgal* and *Itgb1* and in some cases a third round to delete *Itgb7*.

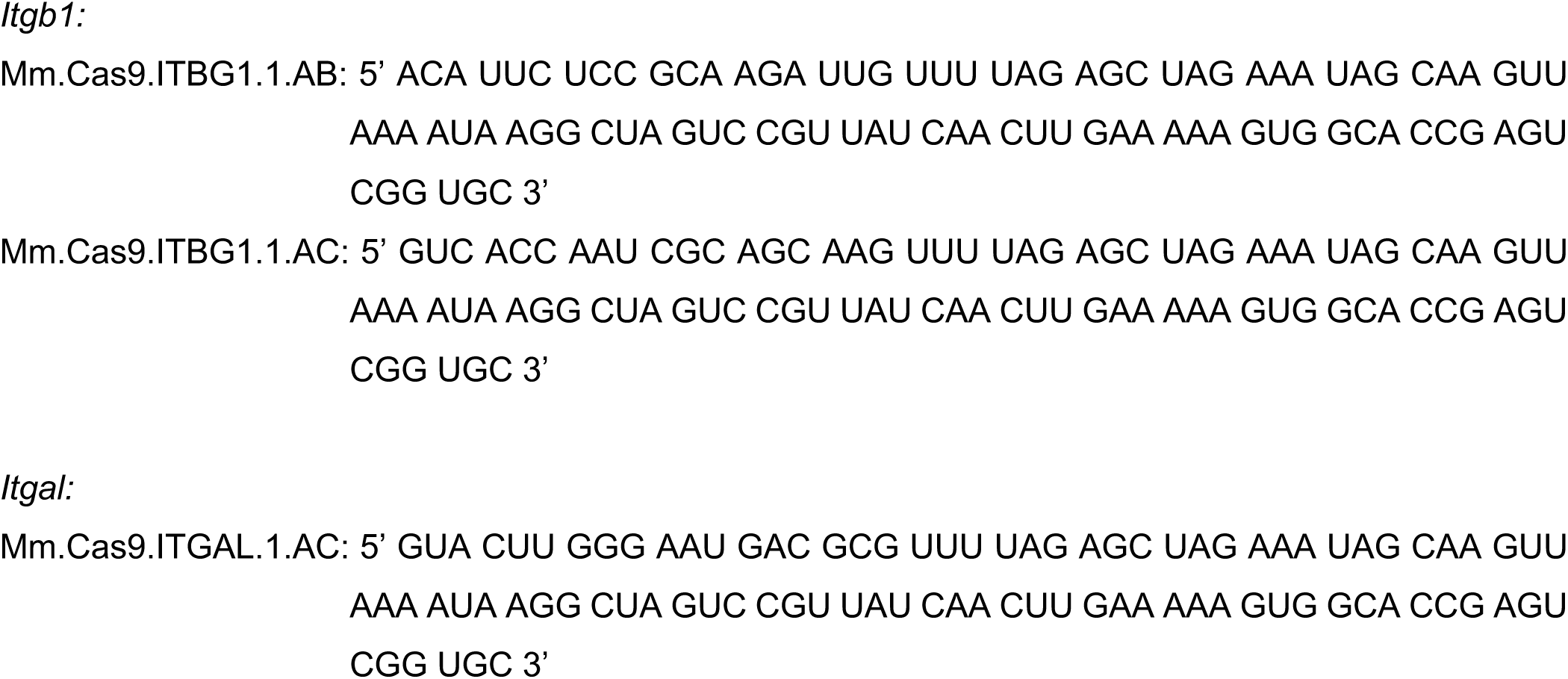
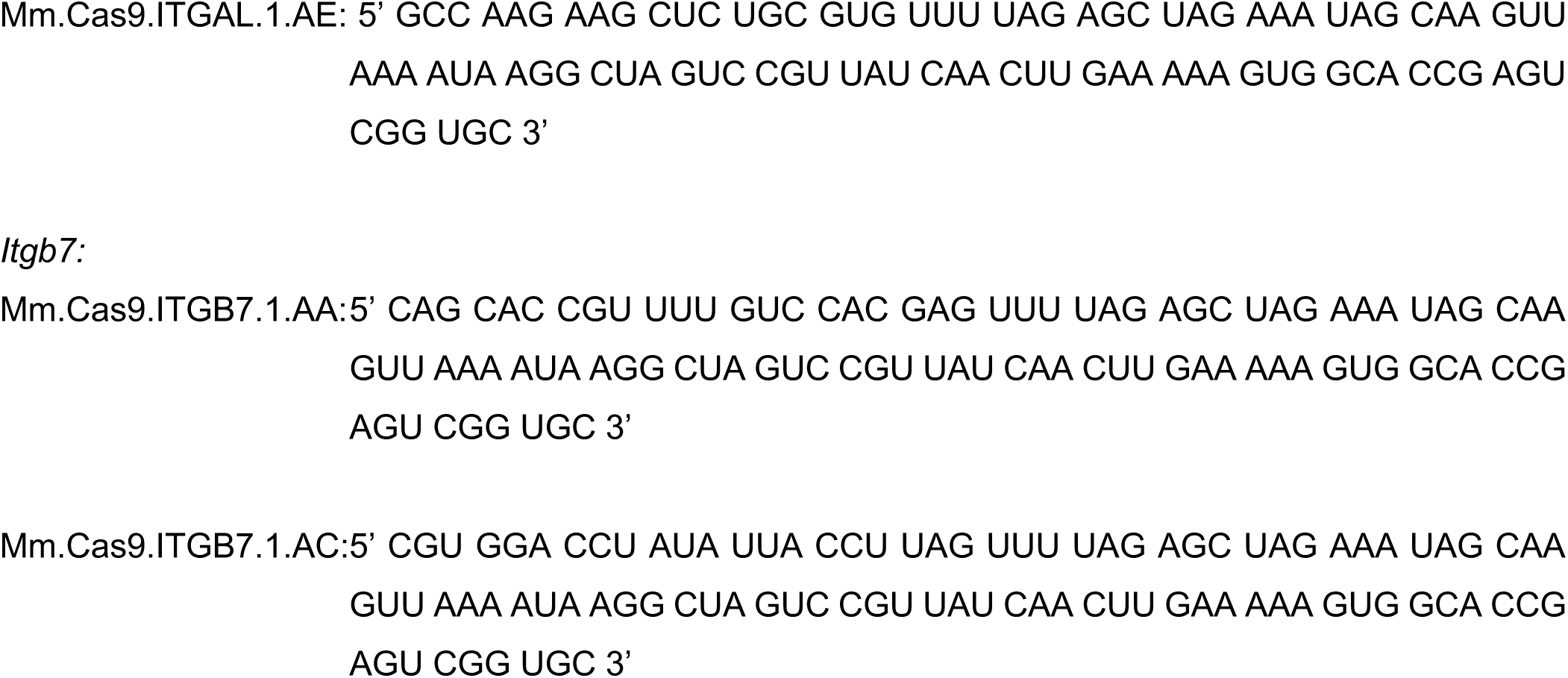

### Tissue isolation and processing for flow cytometry

All mice were euthanized and perfused with 5-10mL PBS. Unless otherwise specified, after isolation, all tissues and cell suspensions were stored in PBS/2% FBS on ice.

*Blood:* collected via cardiac puncture and stored in blood collection tube with K_2_EDTA (BD Microtainer).

*Bone marrow*: right tibia and femur were flushed with PBS/2% FBS.

*Skull bone marrow*: isolated from calvarium by mechanical disruption with glass mortar and pestle. *Spleen and lymph nodes*: mechanical disruption and filtration through a 100um cell strainer with the flat end of a 1mL syringe plunger.

*Central nervous system (CNS):* brain and inside of the dura attached to skull were washed with PBS/2% FBS via p1000 pipette and cells were collected. The wash and whole brain were placed in a 50mL conical tube and pulse-vortexed on low speed for 15 seconds. Brain was collected and mechanically disrupted and filtered through a 100um cell strainer with the flat end of a 1mL syringe plunger. Lymphocytes were purified by centrifugation using a 33% isotonic density gradient (Percoll; GE Healthcare). Red blood cells were depleted using tris-ammonium chloride solution. For later experiments (Fig. 1K,L, 2D-4), we determined that brain parenchyma processing was dispensable: >95% of T-ALL cells were collected with just wash of inside of dura and vortex of whole brain, and ratio was unaffected. All mice in each experiment were processed using the same protocol.

*Cell counting*, single-cell suspensions were counted using a Multisizer 3 (Beckman Coulter) set to detect nuclei between 3.5-10um. Total cell counts from bone marrow were obtained by assuming that cells isolated from tibia and femur of one leg represent 10% of the total bone marrow per mouse (unless otherwise noted). Total cell counts from blood were obtained by assuming that the total amount of blood per mouse is 2mL.

### Immunostaining and flow cytometry

*Immunostaining:* Unless otherwise specified, surface stains were performed on live cells with fluorophore-conjugated antibodies at 1:200 dilution in PBS supplemented with 2% FBS, 2% Normal Mouse Serum (NMS), and 2% Normal Rat Serum (NRS), for 20 minutes on ice and protected from light.

*Antibodies purchased from Biolegend:* anti-B220 (clone RA3-6B2), anti-CD4 (clone GK1.1), anti-CD8 (clone 53-6.7), anti-CD11a (clone M17/4), anti-CD11b (clone M1/70), anti-CD25 (clone PC61), anti-CD31 (clone MEC13.3), anti-CD44 (clone IM7), anti-CD45 (clone 30-F11), anti CD45.1 (clone A20), anti-CD45.2 (clone 104), anti-CD49d (clone R1-2), anti-CD104 (clone 346-11A), anti-β7 (clone FIB504), anti-CX3CR1 (SA011F11), anti-GR1 (clone RB6-8C5), anti-Ter119 (clone TER-119), Streptavidin, mouse IgG2a isotype control (clone MOPC-173) and rat IgG2a, k isotype control (clone RTK2758).

*Antibodies purchased from eBioscience:* anti-CD4 (clone GK1.5), anti-CD29 (clone eBioHMb1-1), anti-Lyve1 (clone ALY7), and Armenian Hamster IgG isotype control (clone eBio299Arm).

*Apoptosis stains*: anti-Active Caspase-3 (clone C92-605) was purchased from BD Biosciences and cells were fixed and permeabilized according to kit instructions (BD Biosciences Cytofix/Cytoperm Cat: BDB554714). Cells were stained at a concentration of 1:100 for 20 minutes at 4C.

*Viability stains:* “LIVE/DEAD™ Fixable Far Red Dead Cell Stain Kit, for 633 or 635 nm excitation” was purchased and used according to kit instructions (ThermoFisher).

*EdU labelling:* “Click-iT™ Plus EdU Alexa Fluor™ 488 Flow Cytometry Assay Kit” and “Click-iT™ Plus EdU Alexa Fluor™ 647 Flow Cytometry Assay Kit” were purchased and cells were stained according to kit instructions (ThermoFisher). DNA quantification was performed by staining with DAPI (Sigma) at 2ug/mL with 5ug/ml RNAse A (Thermo Scientific) for 15 minutes at RT.

*Flow cytometry:* data was acquired using a BD LSRII or Beckman Coulter Cytoflex and analyzed using FlowJo v10.

### Mouse treatments

*For partial irradiation/shielding of CNS:* 1/16” sheet lead (McMaster-Carr) was formed into a half cylinder (6” long, 2” radius) with two semi-circular cut outs in the middle of the long sides (approx. 1-1.5” deep). Four of these half-cylinders were nested to achieve a total thickness of ¼” inch lead and secured with tape. Mice were anesthetized with ketamine/xylazine (100mg/kg /10 mg/kg) and placed on a rigid, irradiator-safe surface with their limbs taped out perpendicular to their body. The lead shielding was placed over the mice, ensuring that their taped limbs were exposed by the semi-circular cut outs, and that all other parts of their body, including the tail, were underneath the lead. Mice were irradiated with 550 cGy using an X-ray irradiator with directional collimator and SnAlCu filter (X-rad320, Precision X-Ray).

*For ablation of meningeal lymphatics,* 5–8-week-old female C57BL/6 mice were injected intraperitoneally with 1*10^12^ PFU of either AAV-VEGFR3(1-4)-Ig (active) or AAV-VEGFR3(4-7)-Ig (control) (provided by Kari Alitalo).

*For intraventricular infusion,* mice were surgically implanted with an intraventricular cannula (P1 Technologies, C315GS-4/SP GUIDE 26GA 38834 4MM Pedestal, 2MM Projection) in the left lateral ventricle (-0.5mm R/C, -1.0mm M/L, -1.8mm D/V relative to bregma). For infusions, >5uL was drawn up into a 33G air-tight Hamilton Syringe (Hamilton, 1705 series) and syringe was mounted in a Remote Infuse/Withdraw Pump 11 Elite Nanomite Programmable Syringe Pump (Harvard Apparatus) mounted on a stereotaxic instrument (Model 940, Kopf). Mice were anesthetized with isoflurane (1.5%-2.5%) or ketamine/xylazine (100mg/kg / 10mg/kg) and mounted via bite bar/nose cone. Syringe needle was advanced into cannula 8.5mm from opening (approx. 0.3-0.5mm into lateral ventricle) and 5uL infusion was conducted at 250nL/min with a 10min rest before needle was withdrawn. *For dye infusion,* NHS-ester-Atto647 (Sigma) was dissolved in DMSO at 10mg/mL (stored at -20C), then further diluted 1:5 in PBS to 2mg/mL for infusions.

*For myeloid cell depletion*, CSF1R inhibitor PLX5622 was formulated at 1200ppm into rodent diet AIN-76A by Research Diets, Inc. (New Brunswick, NJ). Animals received either the PLX5622 formulated diet or a normal chow diet (AIN-76A) a week prior to leukemia induction and for the duration of the experiment (total four weeks). Both PLX5622 and control diet were irradiated and contained red dye.

*For EdU labelling experiments*, mice were administered 100ug EdU (ThermoFisher) in 100uL PBS intraperitoneally at 4h and 2h before euthanasia for a total dose of 200ug EdU.

*For 5-fluorouracil experiments,* 5-fluorouracil (5FU) (Sigma) was dissolved in DMSO to a concentration of 50mg/mL and stored at -20C. For injections, 5FU was diluted 1:10 in PBS to 5mg/mL and mice were injected intraperitoneally at a dose of 40mg/kg. Vehicle-treated mice were injected with 150uL of 10% DMSO/PBS.

*For blocking antibody experiments,* intracerebral ventricular infusions were performed as described above. Anti-CD49d (PS/2; Bio-Xcell) and anti-CD11a (M17/4; Bio-Xcell) were combined at equal quantities for a total of 26ug of each in 5uL. Isotype control mice received 5uL (47ug) of Rat IgG2a, anti-trinitrophenol (clone 2A3; Bio-Xcell). Infusions were performed three days and one day before tissue isolation and analysis.

### Immunofluorescence

*Meninges whole mounts*, adapted from [72]. The top of the skull was removed with surgical scissors. While still attached to the skull, whole-mount meninges were fixed in 2% PFA overnight at 4C. The dura/arachnoid was then peeled from the skullcap. Meninges were incubated in blocking solution [PBS containing 2% normal serum (goat and/or donkey), 3% bovine serum albumin (BSA) (Fisher), anti-CD16/32 (clone 93, eBioscience, 1:1,000), 0.1% Triton X- 100, and 0.05% Tween 20] supplemented with avidin block (Vector Laboratories) for one hour at RT on a shaker; washed (with PBS 3 times for 5 min each at RT); followed by another one-hour block at RT with blocking buffer supplemented with biotin block (Vector Laboratories), and washed again. Primary antibodies were applied at 1:200 dilution in staining buffer [PBS containing 3% BSA and 0.5% Triton X-100] and meninges were incubated overnight at 4C. Meninges were washed then incubated in appropriate dilutions of secondary antibodies in staining buffer for 2 hours at RT or overnight at 4C. Meninges were washed, then wet mounted on a Superfrost Plus glass slide, and allowed to dry for 10-15 minutes. Fluoromount G and a coverslip were added and slides were allowed to fully dry at RT (16h) before edges were sealed with clear nail polish.

*Primary antibodies*: anti-Lyve-1 biotinylated (clone ALY7; eBioscience), anti-CD31-PE (clone MEC13.3; Biolegend), anti-GFP (ab13970; Abcam), anti-RFP (600-401-379, Rockland).

*Secondary antibodies,* Streptavidin-BV421 1:500 (Biolegend), Alexa Fluor 488 AffiniPure Goat Anti-Chicken IgY (IgG) (H+L) 1:1000 (Jackson ImmunoResearch), Alexa Fluor 647 AffiniPure Donkey Anti-Rabbit IgG (H+L) 1:1000 (Jackson ImmunoResearch), Cy3 Affinipure Donkey anti-Rabbit IgG (H+L) 1:1000 (Jackson ImmunoResearch).

Images were acquired using confocal microscope (LSM710 (Carl Zeiss) or Stellaris 8 (Leica)) with a 10x dry or 20x dry objective using ZEN (Zeiss) or LasX (Leica) software. Images were processed and analyzed using ImageJ (version 1.53; National Institutes of Health) and FIJI (version 2.14.0, Schindelin et al, 2012).

### RNA extraction and sequencing

*For T-ALL cell sequencing:* DKO, WT, CRISPR-DKO and CRISPR-WT (both CD45.2) cells were isolated from the CNS and spleen of 1:1 DKO:WT co-transfer mice (congenically-marked,CD45.1/.2) at mid-stage disease, stained for the presence of the integrins and CD45.1 and CD45.2, and sorted using a BD FACS Aria.

*For fibroblast sequencing:* Dural meninges were isolated from naïve wild-type mice as described in [26] and placed immediately in microcentrifuge tubes of digestion buffer containing collagenase D (1mg/ml, MilliporeSigma), DNAse I (50ug/mL, Roche), and Dispase (5 U/mL, Roche) in serum-free, high-glucose DMEM (Invitrogen) (modified from [43]) and stored on ice. Tissues in microtubes were placed on rotator for 2h at 37C. Suspension was agitated by pipetting until fully dissociated and filtered through 40um cell strainer. Single cell suspension was pelleted at 400g for 4 minutes and seeded on 6-well tissue culture treated plates in meningeal fibroblast media (high-glucose DMEM supplemented with 15% FBS (Cytiva) and 100 IU/mL penicillin and 100 ug/mL streptomycin (Gibco or Corning)). 16h after seeding, media and non-attached cell fraction was aspirated, leaving fibroblasts behind, and media was replaced and changed every 2-3 days. Meningeal fibroblasts were cultured for 10-14 days before being lifted from the plate with Trypsin/EDTA and snap-frozen in Trizol for RNA extraction.

Total RNA was extracted using RNeasy Plus Micro Kit (Qiagen) and purified with RNAse-Free DNAse (Qiagen). Total RNA was quantified on an Agilent BioAnalyzer via the Pico kit (Agilent). SMART-Seq HT Kit (Takara) was used to construct cDNA and libraries. cDNA was constructed using 1ng input of RNA and amplified with 12 cycles. cDNA product was purified with DNA AMPure beads (Beckman-Coulter) and quantified via Agilent Quant-IT. 300pg of each cDNA sample was used to construct libraries via Nextera XT DNA library preparation kit. 12 PCR cycles were used to amplify the libraries and to add the barcodes. Libraries were then cleaned, quantified, and pooled for sequencing on an S1 (100 cycles) flow cell on Illumina NovaSeq sequencer.

Sequencing reads were mapped to the reference genome (mm10) using the STAR aligner (v2.5.0c) [73]. Alignments were guided by a Gene Transfer Format (GTF) file. The mean read insert sizes and their standard deviations were calculated using Picard tools (v.1.126) (http://broadinstitute.github.io/picard). The read count tables were generated using HTSeq (v0.6.0) [74], normalized based on their library size factors using DEseq2 [75], and differential expression analysis was performed. The Read Per Million (RPM) normalized BigWig files were generated using BEDTools (v2.17.0) [76] and bedGraphToBigWig tool (v4). All the downstream statistical analyses and generating plots were performed in R environment (v3.1.1) (https://www.r-project.org/).

Nichenet analysis was performed according to [48, 49] and using the code graciously provided on https://github.com/saeyslab/nichenetr/blob/master/vignettes/ligand_activity_geneset.md. The list of diferentially expressed genes was identified by FDR <0.1 from the DESeq2 analysis performed on the WT and DKO T-ALL isolated from the CNS (described above). The mouse reference libraries were used for analysis.

Packages used to generate plots were TidyVerse [77], ggplot, and circlize [78].

### Statistical analysis

Statistical analysis (excluding RNA-seq experiments) was conducted using the PRISM program (GraphPad). Experiments comparing the total number of WT to the total number of DKO T-ALL cells, and experiments comparing the ratio of DKO:WT between groups, were analyzed using a two-tailed, unpaired t-test with Welch’s correction. Experiments comparing the survival or proliferation of WT to DKO T-ALL cells were analyzed using a two-tailed, paired t-test. For data displayed on a log-scale, log_10_-transformed data was used to calculate significance. For data presented as percentages, logit-transformed data was used to calculate significance. Significance was defined as ∗p < 0.05, ∗∗p < 0.01, ∗∗∗p < 0.001, ∗∗∗∗p < 0.0001.

## Online supplemental material

S. Fig. 1 shows how T-ALL is characterized by flow cytometry. S. Fig. 2 shows gene expression by T-ALL cells assessed by RNA sequencing. S. Fig. 3 shows depletion of meningeal lymphatics by the VEGF-trap and depletion of macrophages by CSF1R inhibition. S. Fig. 4 shows depletion of CNS T-ALL by 5FU and lack of depletion of T-ALL by antibodies to a4 and aL integrins alone.

## Data availability

RNA sequencing expression data included in this study have been deposited in the NCBI Gene Expression Omnibus (GEO), accession number GSE318579 (reviewer access code obmdyoqevzivxib). Additional data not in the article itself are available upon reasonable request from the corresponding author.

## Acknowledgements

We thank members of the Schwab laboratory for helpful discussions throughout the duration of the project, and Eva Hernando for her insightful suggestions and review of the manuscript. We thank Kari Alitalo (University of Helsinki) for the VEGFR3 and control AAV and the K14-VEGFR3-Ig mice, and Iannis Aifantis (NYU School of Medicine) for the *Notch*Δ*E*-GFP/mCherry vectors. We are grateful to the NYU Langone Health Genome Technology Center and Applied Bioinformatics Laboratories, especially Alireza Khodadadi-Jamayran, for assistance with RNA sequencing and bioinformatic analysis. We would like to thank the NYU Langone Health Division of Comparative Medicine animal technicians, especially Abdoul Fall, for their assistance.

This work was funded by a Blood Cancer Discoveries Grant from Blood Cancer United, the Mark Foundation for Cancer Research, and the Paul G. Allen Frontiers Group (SRS); a pilot project grant from the NYU Langone Center for Blood Cancers (SRS); a Vilcek MSTP Scholarship (SYL); National Institutes of Health grant T32AI100853 (SYL, CC, BSS); National Institutes of Health grant UL1RR029893 (CC); National Institutes of Health grant F30CA298305 (HC); National Institutes of Health grant F31AI154793 (MO); National Institutes of Health grant T32GM136542 (BSS); National Institutes of Health grant R01HD088411 (RCF); National Institutes of Health grant U19NS107616 (RCF); and National Institutes of Health grant P30CA016087.

Author contributions are as follows. Conceptualization: SYL, CC, SRS. Methodology: MO, KAM, AYC, JKS, RCF, MC. Investigation: SYL, CC, HC, BSS, JBV. Formal analysis: SYL, CC. Writing, original draft: SYL, SRS. Writing, review & editing: SYL, CC, HC, BSS, SRS. Supervision: SRS.

The authors have no conflicting financial interests.

## References

1. Cronin, K.A., et al., Annual report to the nation on the status of cancer, part 1: National cancer statistics. Cancer, 2022. 128(24): p. 4251–4284.

2. Inaba, H. and C.H. Pui, Advances in the Diagnosis and Treatment of Pediatric Acute Lymphoblastic Leukemia. J Clin Med, 2021. 10(9).

3. Thastrup, M., et al., Central nervous system involvement in childhood acute lymphoblastic leukemia: challenges and solutions. Leukemia, 2022. 36(12): p. 2751–2768.

4. Frishman-Levy, L. and S. Izraeli, Advances in understanding the pathogenesis of CNS acute lymphoblastic leukaemia and potential for therapy. Br J Haematol, 2017. 176(2): p. 157–167.

5. Hunger, S.P. and C.G. Mullighan, Acute Lymphoblastic Leukemia in Children. N Engl J Med, 2015. 373(16): p. 1541–52.

6. Pui, C.H. and S.C. Howard, Current management and challenges of malignant disease in the CNS in paediatric leukaemia. Lancet Oncol, 2008. 9(3): p. 257–68.

7. Engelhardt, B. and R.M. Ransohoff, Capture, crawl, cross: the T cell code to breach the blood-brain barriers. Trends Immunol, 2012. 33(12): p. 579–89.

8. Yednock, T.A., et al., Prevention of experimental autoimmune encephalomyelitis by antibodies against alpha 4 beta 1 integrin. Nature, 1992. 356(6364): p. 63-6.

9. Baron, J.L., et al., Surface expression of alpha 4 integrin by CD4 T cells is required for their entry into brain parenchyma. J Exp Med, 1993. 177(1): p. 57–68.

10. Rudick, R., et al., Natalizumab: bench to bedside and beyond. JAMA Neurol, 2013. 70(2): p. 172–82.

11. Yao, H., et al., Leukaemia hijacks a neural mechanism to invade the central nervous system. Nature, 2018. 560(7716): p. 55-60.

12. Pitt, L.A., et al., CXCL12-Producing Vascular Endothelial Niches Control Acute T Cell Leukemia Maintenance. Cancer Cell, 2015. 27(6): p. 755–68.

13. Liu, Y., et al., The genomic landscape of pediatric and young adult T-lineage acute lymphoblastic leukemia. Nat Genet, 2017. 49(8): p. 1211–1218.

14. Weng, A.P., et al., Activating mutations of NOTCH1 in human T cell acute lymphoblastic leukemia. Science, 2004. 306(5694): p. 269-71.

15. Pear, W.S., et al., Exclusive development of T cell neoplasms in mice transplanted with bone marrow expressing activated Notch alleles. J Exp Med, 1996. 183(5): p. 2283–91.

16. Ding, Z.M., et al., Relative contribution of LFA-1 and Mac-1 to neutrophil adhesion and migration. J Immunol, 1999. 163(9): p. 5029–38.

17. Raghavan, S., et al., Conditional ablation of beta1 integrin in skin. Severe defects in epidermal proliferation, basement membrane formation, and hair follicle invagination. J Cell Biol, 2000. 150(5): p. 1149–60.

18. Luo, B.H., C.V. Carman, and T.A. Springer, Structural basis of integrin regulation and signaling. Annual review of immunology, 2007. 25: p. 619–47.

19. Price, R.A. and W.W. Johnson, The central nervous system in childhood leukemia. I. The arachnoid. Cancer, 1973. 31(3): p. 520–33.

20. Zohren, F., et al., The monoclonal anti-VLA-4 antibody natalizumab mobilizes CD34+ hematopoietic progenitor cells in humans. Blood, 2008. 111(7): p. 3893–5.

21. Papayannopoulou, T., et al., The VLA4/VCAM-1 adhesion pathway defines contrasting mechanisms of lodgement of transplanted murine hemopoietic progenitors between bone marrow and spleen. Proc Natl Acad Sci U S A, 1995. 92(21): p. 9647–51.

22. Bonig, H., et al., Increased numbers of circulating hematopoietic stem/progenitor cells are chronically maintained in patients treated with the CD49d blocking antibody natalizumab. Blood, 2008. 111(7): p. 3439–41.

23. Berlin, C., et al., Alpha 4 beta 7 integrin mediates lymphocyte binding to the mucosal vascular addressin MAdCAM-1. Cell, 1993. 74(1): p. 185–95.

24. Meeker, R.B., et al., Cell trafficking through the choroid plexus. Cell Adh Migr, 2012. 6(5): p. 390–6.

25. Rossi, B., et al., Alpha4 beta7 integrin controls Th17 cell trafficking in the spinal cord leptomeninges during experimental autoimmune encephalomyelitis. Front Immunol, 2023. 14: p. 1071553.

26. Louveau, A., et al., Structural and functional features of central nervous system lymphatic vessels. Nature, 2015. 523(7560): p. 337-41.

27. Aspelund, A., et al., A dural lymphatic vascular system that drains brain interstitial fluid and macromolecules. J Exp Med, 2015. 212(7): p. 991–9.

28. Teijeira, A., et al., T Cell Migration from Inflamed Skin to Draining Lymph Nodes Requires Intralymphatic Crawling Supported by ICAM-1/LFA-1 Interactions. Cell Rep, 2017. 18(4): p. 857–865.

29. Hunter, M.C., A. Teijeira, and C. Halin, T Cell Trafficking through Lymphatic Vessels. Front Immunol, 2016. 7: p. 613.

30. Antila, S., et al., Development and plasticity of meningeal lymphatic vessels. J Exp Med, 2017. 214(12): p. 3645–3667.

31. Dustin, M.L., The immunological synapse. Cancer Immunol Res, 2014. 2(11): p. 1023–33.

32. Zamora, A.E., et al., Pediatric patients with acute lymphoblastic leukemia generate abundant and functional neoantigen-specific CD8. Sci Transl Med, 2019. 11(498).

33. Li, Y., et al., Impact of T-cell immunity on chemotherapy response in childhood acute lymphoblastic leukemia. Blood, 2022. 140(13): p. 1507–1521.

34. Lyu, A., et al., Tumor-associated myeloid cells provide critical support for T-ALL. Blood, 2020. 136(16): p. 1837–1850.

35. Lyu, A., et al., Integrin signaling is critical for myeloid-mediated support of T-cell acute lymphoblastic leukemia. Nat Commun, 2023. 14(1): p. 6270.

36. Spangenberg, E., et al., Sustained microglial depletion with CSF1R inhibitor impairs parenchymal plaque development in an Alzheimer’s disease model. Nat Commun, 2019. 10(1): p. 3758.

37. Lei, F., et al., CSF1R inhibition by a small-molecule inhibitor is not microglia specific; affecting hematopoiesis and the function of macrophages. Proc Natl Acad Sci U S A, 2020. 117(38): p. 23336–23338.

38. Montilla, A., et al., Microglia and meningeal macrophages depletion delays the onset of experimental autoimmune encephalomyelitis. Cell Death Dis, 2023. 14(1): p. 16.

39. Humphrey, R.S., et al., Tumor-associated myeloid cells support murine T-ALL in the central nervous system via integrin signaling. Blood Neoplasia, 2025. 2(3): p. 100095.

40. Schlager, C., et al., Effector T-cell trafficking between the leptomeninges and the cerebrospinal fluid. Nature, 2016. 530(7590): p. 349-53.

41. Jonart, L.M., et al., Disrupting the leukemia niche in the central nervous system attenuates leukemia chemoresistance. Haematologica, 2020. 105(8): p. 2130–2140.

42. Derk, J., et al., Living on the Edge of the CNS: Meninges Cell Diversity in Health and Disease. Front Cell Neurosci, 2021. 15: p. 703944.

43. Remsik, J., et al., Characterization, isolation, and in vitro culture of leptomeningeal fibroblasts. J Neuroimmunol, 2021. 361: p. 577727.

44. DeSisto, J., et al., Single-Cell Transcriptomic Analyses of the Developing Meninges Reveal Meningeal Fibroblast Diversity and Function. Dev Cell, 2020. 54(1): p. 43–59.e4.

45. Davidson, S., et al., Fibroblasts as immune regulators in infection, inflammation and cancer. Nat Rev Immunol, 2021. 21(11): p. 704–717.

46. Buechler, M.B. and S.J. Turley, A short field guide to fibroblast function in immunity. Semin Immunol, 2018. 35: p. 48–58.

47. Ramaglia, V., et al., Stromal Cell-Mediated Coordination of Immune Cell Recruitment, Retention, and Function in Brain-Adjacent Regions. J Immunol, 2021. 206(2): p. 282–291.

48. Browaeys, R., W. Saelens, and Y. Saeys, NicheNet: modeling intercellular communication by linking ligands to target genes. Nat Methods, 2020. 17(2): p. 159–162.

49. Bonnardel, J., et al., Stellate Cells, Hepatocytes, and Endothelial Cells Imprint the Kupffer Cell Identity on Monocytes Colonizing the Liver Macrophage Niche. Immunity, 2019. 51(4): p. 638–654.e9.

50. Myers, C.E., et al., Pharmacokinetics of the fluoropyrimidines: implications for their clinical use. Cancer Treat Rev, 1976. 3(3): p. 175–83.

51. Bourke, R.S., et al., Kinetics of entry and distribution of 5-fluorouracil in cerebrospinal fluid and brain following intravenous injection in a primate. Cancer Res, 1973. 33(7): p. 1735–46.

52. Levin, V.A., M. Chadwick, and A.D. Little, Distribution of 5-fluorouracil-2- 14 C and its metabolites in a murine glioma. J Natl Cancer Inst, 1972. 49(6): p. 1577–84.

53. Clarkson, B., et al., THE PHYSIOLOGIC DISPOSITION OF 5-FLUOROURACIL AND 5-FLUORO-2’-DEOXYURIDINE IN MAN. Clin Pharmacol Ther, 1964. 5: p. 581–610.

54. Rustenhoven, J., et al., Functional characterization of the dural sinuses as a neuroimmune interface. Cell, 2021. 184(4): p. 1000–1016.e27.

55. Rua, R. and D.B. McGavern, Advances in Meningeal Immunity. Trends Mol Med, 2018. 24(6): p. 542–559.

56. Ko, S.Y., et al., High CXCR4 and low VLA-4 expression predicts poor survival in adults with acute lymphoblastic leukemia. Leuk Res, 2014. 38(1): p. 65–70.

57. Cleaver, A.L., et al., Gene-based outcome prediction in multiple cohorts of pediatric T-cell acute lymphoblastic leukemia: a Children’s Oncology Group study. Mol Cancer, 2010. 9: p. 105.

58. Hsieh, Y.T., et al., Integrin alpha4 blockade sensitizes drug resistant pre-B acute lymphoblastic leukemia to chemotherapy. Blood, 2013. 121(10): p. 1814–8.

59. Shalapour, S., et al., High VLA-4 expression is associated with adverse outcome and distinct gene expression changes in childhood B-cell precursor acute lymphoblastic leukemia at first relapse. Haematologica, 2011. 96(11): p. 1627–35.

60. Ruan, Y., et al., In vitro and in vivo effects of AVA4746, a novel competitive antagonist of the ligand binding of VLA-4, in B-cell acute lymphoblastic leukemia. Exp Ther Med, 2022. 23(1): p. 47.

61. Berrazouane, S., et al., Beta1 integrin blockade overcomes doxorubicin resistance in human T-cell acute lymphoblastic leukemia. Cell Death Dis, 2019. 10(5): p. 357.

62. Hsieh, Y.T., et al., Effects of the small-molecule inhibitor of integrin α4, TBC3486, on pre-B-ALL cells. Leukemia, 2014. 28(10): p. 2101–4.

63. Matsunaga, T., et al., Interaction between leukemic-cell VLA-4 and stromal fibronectin is a decisive factor for minimal residual disease of acute myelogenous leukemia. Nat Med, 2003. 9(9): p. 1158–65.

64. Winter, S.S., et al., Enhanced T-lineage acute lymphoblastic leukaemia cell survival on bone marrow stroma requires involvement of LFA-1 and ICAM-1. Br J Haematol, 2001. 115(4): p. 862–71.

65. Guo, L., et al., Proteomic analysis of cerebrospinal fluid in pediatric acute lymphoblastic leukemia patients: a pilot study. Onco Targets Ther, 2019. 12: p. 3859–3868.

66. Enlund, S., et al., The CNS microenvironment promotes leukemia cell survival by disrupting tumor suppression and cell cycle regulation in pediatric T-cell acute lymphoblastic leukemia. Exp Cell Res, 2024. 437(2): p. 114015.

