## Supplemental Figures for "Integrin-deficient T cell leukemia accumulates in the central nervous system"

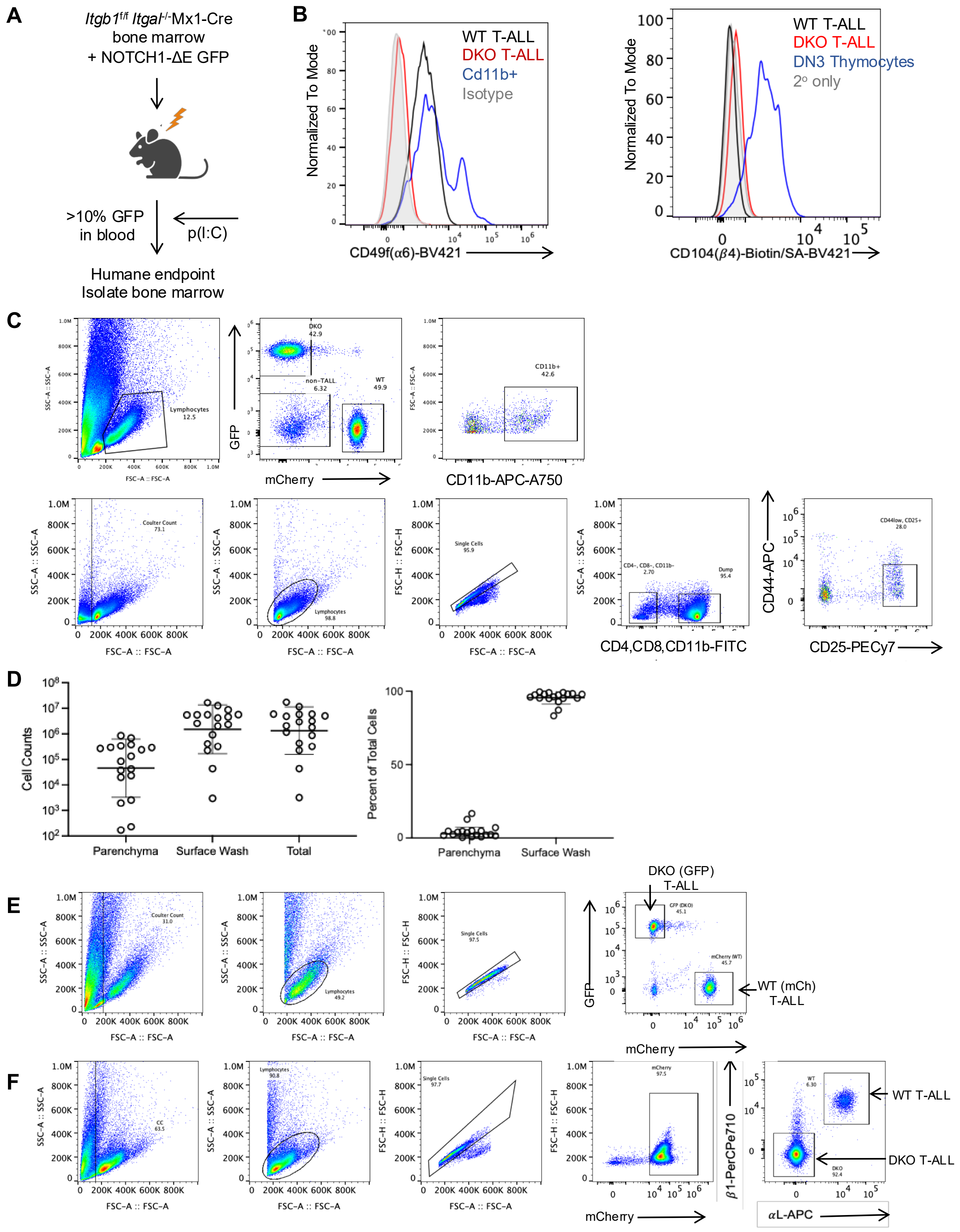


**Supplemental Figure 1: Generation and characterization of WT and DKO T-ALL**

**(A)** Generation of T-ALL cell lines. To generate DKO T-ALL, bone marrow progenitor cells from *Itgal*$^{-/-}$ *Itgb1*$^{f/f}$Mx1-Cre+ mice were transduced with a vector encoding constitutively active NOTCH1 (Notch1-ΔE) and GFP. These cells were transferred intravenously to a lethally irradiated WT recipient (with WT support bone marrow). When GFP+ leukemia cells reached >10% of white blood cells, the mice were treated with the strong interferon inducer polyinosinic-polycytidylic acid (p(I:C)) to activate the Mx1 promoter, induce Cre expression, and cause *Itgb1* excision. The mice were euthanized at a humane endpoint, and GFP+ bone marrow cells were frozen. These cells were used in subsequent transfer experiments. WT control T-ALL was generated similarly, except that the vector encoded mCherry rather than GFP. **(B)** Surface expression of integrin $\alpha$6 (left) or $\beta$4 (right) on WT T-ALL (black) and DKO T-ALL (red) from the CNS, compared to positive control (blue) (CD11b+ cells in the CNS, left; DN3 thymocytes, right) or negative control (grey) (isotype, left; secondary only, right). **(C**) Gating strategy for CD11b+ cells from the CNS (mCherry-, GFP-, Cd11b+) and DN3 thymocytes (CD4-, CD8-, CD11b-, CD44-, CD25+). **(D)** Total number of cells (left) or relative percentage of total cells (right) isolated from the CNS of mice either from parenchyma (by Percoll gradient) or by surface wash (n = 18). Data compiled from five experiments, including both WT:DKO T-ALL and CRISPR-WT:CRISPR-DKO T-ALL co-transfers. **(E)** Gating strategy for identification of DKO (GFP) and WT (mCherry) T-ALL from the CNS. **(F)** Gating strategy for identification of CRISPR-DKO and CRISPR-WT T-ALL from the CNS by surface expression of $\alpha$L and $\beta$1 integrins.

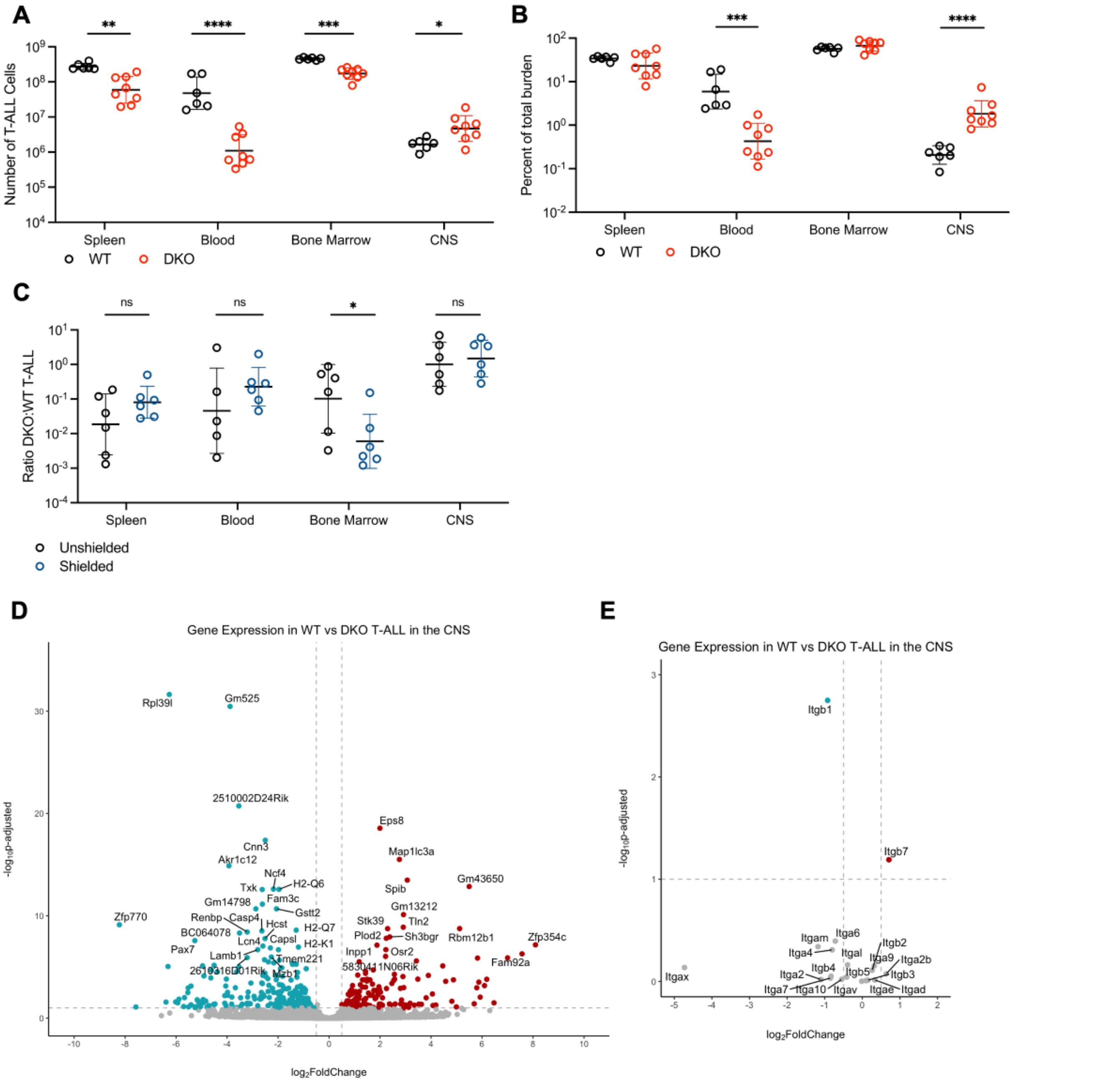


**Supplemental Figure 2: Transcriptomic analysis of integrin-deficient T-ALL**

**(A,B)** Sub-lethally irradiated WT mice received either WT or DKO T-ALL i.v. Number **(A)** or relative disease burden **(B)** of DKO (red) or WT (black) T-ALL cells found in the indicated tissues at a humane endpoint. Data compiled from 2 experiments; n = 8 DKO, n = 6 WT.

(**C**) WT mice received 1:1 DKO:WT T-ALL i.v. after either whole-body sub-lethal irradiation ("unshielded") or sub-lethal irradiation of only the limbs ("shielded"). Ratio of DKO:WT T-ALL in the indicated tissues at mid-stage disease. Data compiled from 2 experiments; n = 6. Statistics calculated on log-transformed data using unpaired t-test with Welch's correction. *$p \leq 0.05$, **$p \leq 0.01$, ***$p \leq 0.001$, ****$p \leq 0.0001$

**(D,E)** RNA-Sequencing was performed on CRISPR-WT and CRISPR-DKO T-ALL cells isolated from the CNS of mice at mid-stage disease that had received 1:1 DKO:WT T-ALL. Volcano plots of **(D)** all differentially expressed genes or **(E)** integrin-family genes. (Right, red) genes that are upregulated in the DKO T-ALL; (Left, blue) genes that are downregulated in the DKO T-ALL. Vertical lines represent log2FoldChange = -0.5, 0.5; horizontal line represents FDR = 0.1. Residual *Itgal* and *Itgb1* expression likely reflects truncated transcripts, as no surface protein remains. Average of *n* = 4 biological replicates. Statistics calculated using DESeq2 and corrected for multiple comparisons.

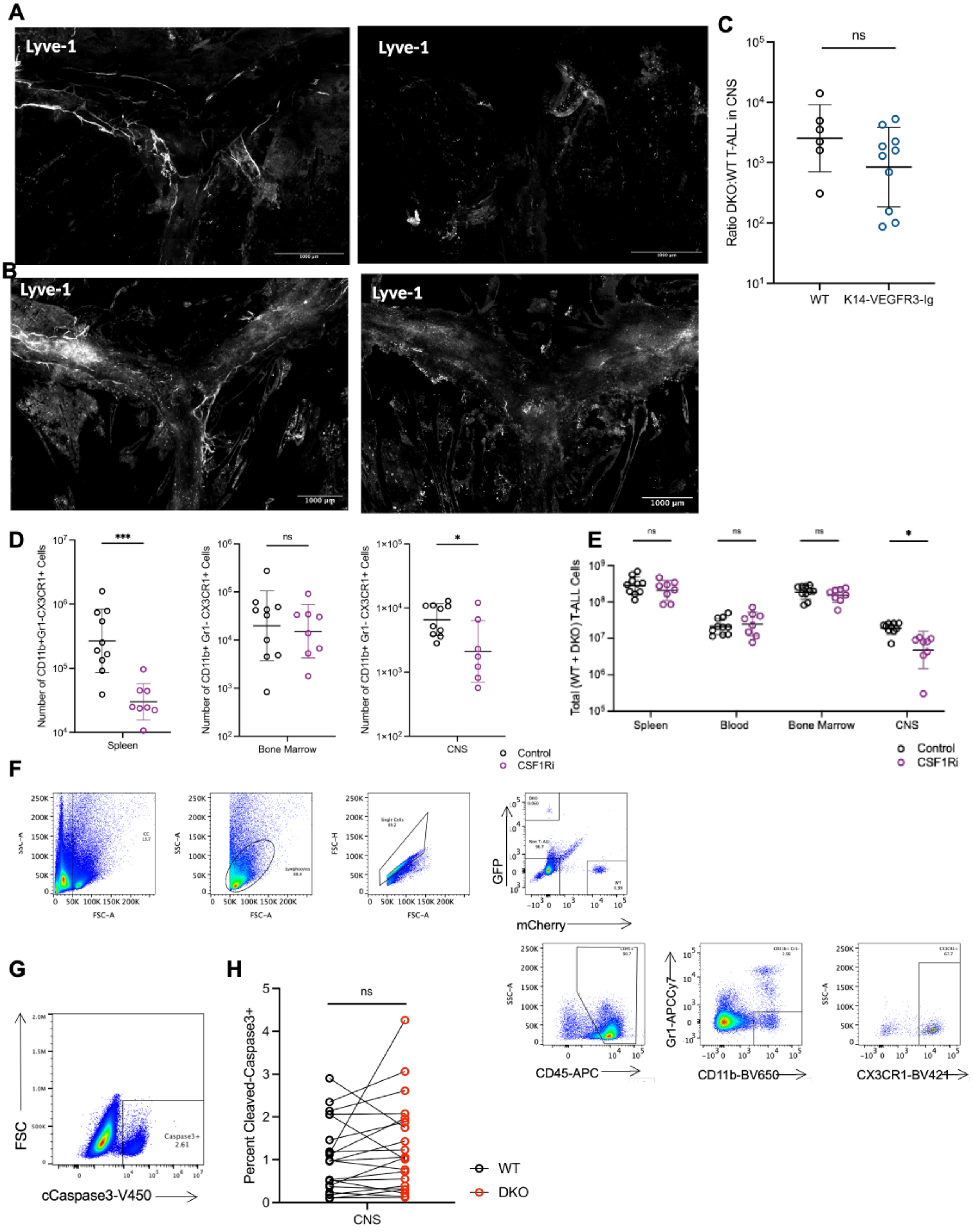


**Supplemental Figure 3: Loss of dorsal meningeal lymphatics and CNS macrophages.**

**(A)** Image of whole-mounted dural meninges from a mouse injected with control AAV (left; representative of *n* = 6 in 2 experiments) or VEGF-trap AAV (right; representative of *n* = 6 in 2 experiments) at end-stage

T-ALL disease. White, anti-Lyve-1; scale bars, 1000$\mu$m. **(B)** Image of whole-mounted dural meninges from a wild-type littermate control (left; representative of *n* = 6 in 2 experiments) or K14-VEGFR3-Ig transgenic mouse (right; representative of *n* = 10 in 2 experiments) at end-stage T-ALL disease. White, anti-Lyve-1; scale bars, 1000$\mu$m. **(C)** Sub-lethally irradiated K14-VEGFR3-Ig transgenic mice (*n* = 10) or wild-type littermate controls (*n* = 6) received 1:1 DKO:WT T-ALL i.v., and were analyzed at end-stage T-ALL disease. Data compiled from 2 experiments. **(D)** The number of tissue-resident macrophages present in the indicated tissues of 1:1 DKO:WT T-ALL mice after four weeks of treatment with either vehicle (black; *n* = 9) or CSF1Ri (PLX5622) diet (purple; *n* = 8). Data compiled from 2 experiments. **(E)** The number of total T-ALL cells (DKO + WT) present in the indicated tissues of 1:1 DKO:WT T-ALL mice after four weeks of treatment with either vehicle (black, *n* = 9) or CSF1Ri (PLX5622) diet (purple; *n* = 8). Data compiled from 2 experiments. **(F)** Gating strategy for identifying tissue-resident macrophages (CD45hi, CD11b+, Gr1-, CX3CR1+). Shown: bone marrow. **(G)** Detection of cleaved caspase 3 in T-ALL cells by flow cytometry. **(H)** Percent cleaved-caspase 3+ WT (black) or DKO (red) T-ALL cells isolated from the CNS of 1:1 DKO:WT T-ALL mice at mid-stage disease (data compiled from 12 experiments; *n* = 21). (C, D, E) Statistics calculated on log-transformed data using unpaired t-test with Welch's correction. (H) Statistics calculated on logit-transformed data using paired t-test.

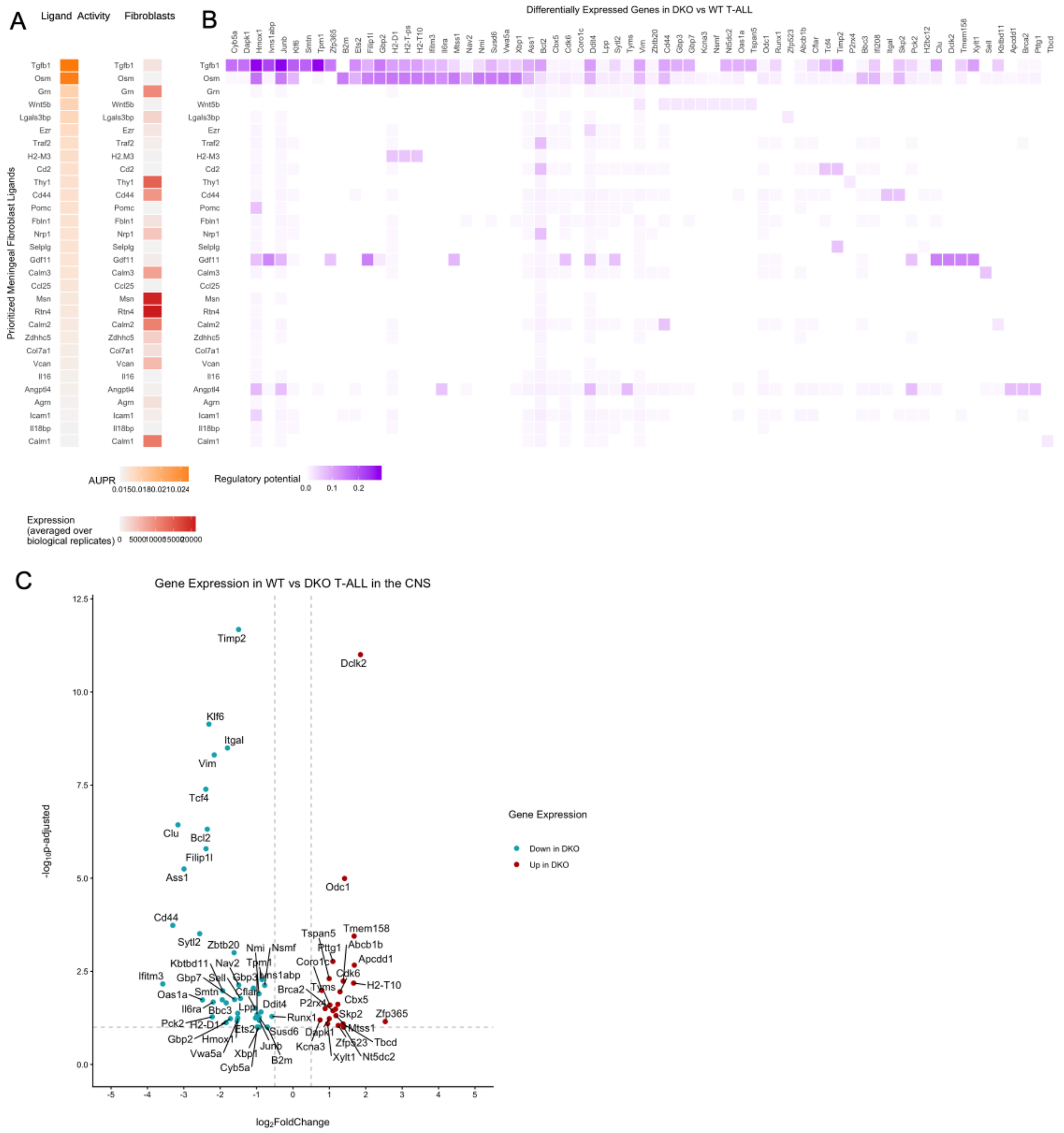


**Supplemental Figure 4: NicheNet analysis of T-ALL and meningeal fibroblasts.**

**(A, B)** NicheNet analysis was performed with significantly differentially expressed genes between DKO and WT T-ALL isolated from the CNS at mid-stage disease (FDR <0.1) (n = 9, 5 DKO:WT co-transfers and 4 CRISPR-WT:CRISPR-DKO co-transfers). These differentially expressed genes were matched to potential ligands expressed by meningeal fibroblasts from the dural meninges of mice with T-ALL at a comparable disease stage expanded in culture (n = 3). **(A)** Prioritized meningeal fibroblast ligands as identified by NicheNet analysis based on how well they fit the modeled ligand-signaling matrix (orange = AUPR, Area Under the Precision-Recall curve, a quantified analysis of how many differentially expressed genes would be affected by downstream signaling from each ligand) and their expression levels in the meningeal fibroblasts (red = expression averaged over biological replicates). **(B)** Matrix of prioritized meningeal fibroblast ligands (y-axis) compared to their downstream targets in the differentially expressed genes between WT and DKO T-ALL (purple = regulatory potential, the likelihood that the differential expression of each gene in the T-ALL cells (x axis) is due to signaling from each ligand (y axis)). **(C)** Volcano plot of the differentially expressed target genes in the WT and DKO T-ALL cells with the highest regulatory potential, showing expression either UP in the DKO (red) or DOWN in the DKO (blue).

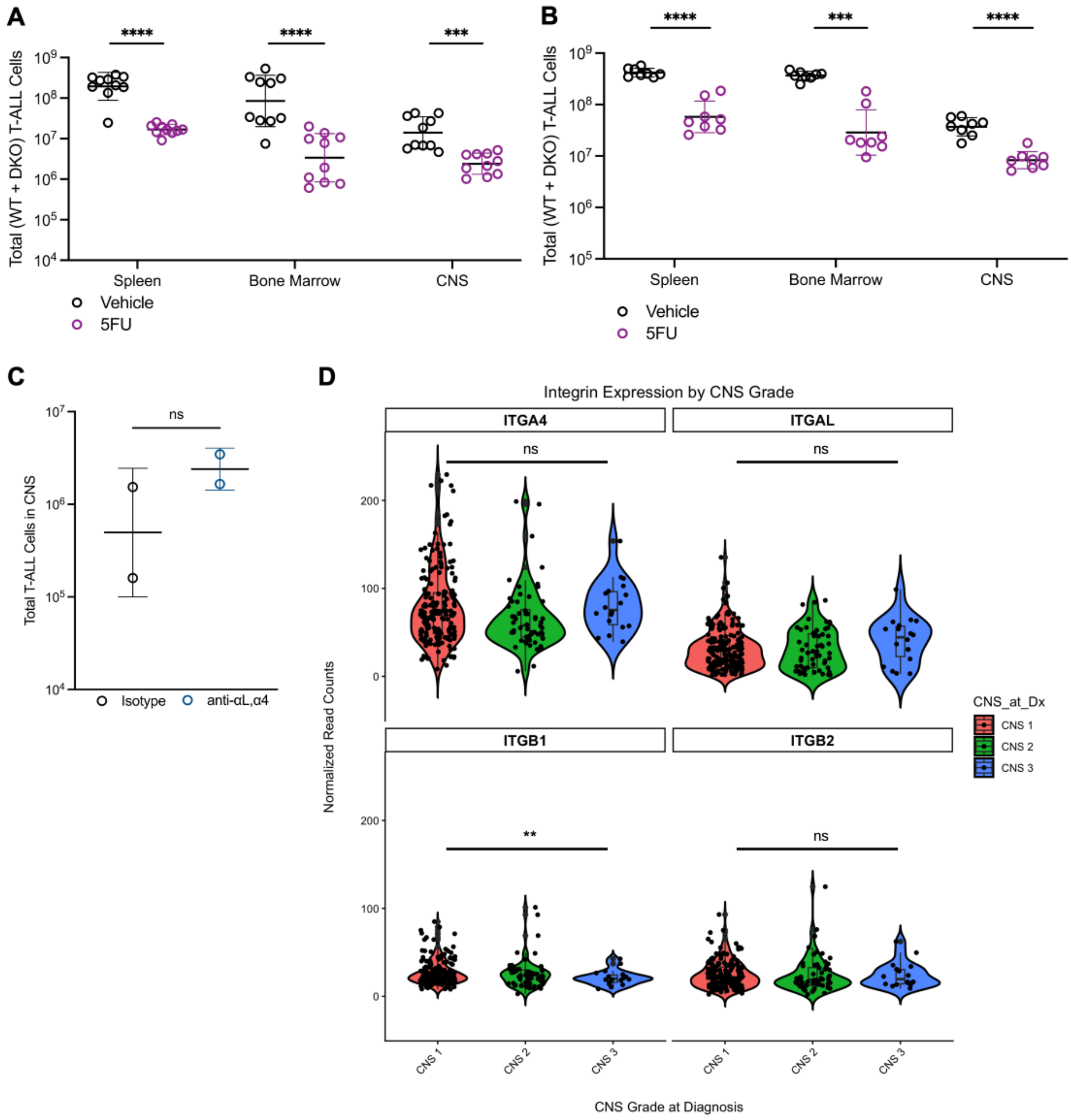


**Supplemental Figure 5: Depletion of T-ALL with 5FU and integrin blockade**

**(A)** Sub-lethally irradiated WT mice received 1:1 DKO:WT T-ALL i.v.; were treated i.p. with 40mg/kg 5-fluorouracil (5-FU) daily for three days at mid-stage disease (12-14 days after T-ALL transfer); and were analyzed 24h after the last dose of 5-FU. Total number of T-ALL cells (WT + DKO) in the indicated tissues after treatment with vehicle (black; *n* = 10) or 5-FU (purple; *n* = 10). Data compiled from 2 experiments. **(B)** As in (A) but for the CRISPR-WT and CRISPR-DKO pair of T-ALL lines. Vehicle-treated, black (*n* = 8); 5FU-treated, purple (*n* = 8). Data compiled from 2 experiments. Statistics calculated on log-transformed data using unpaired t-test with Welch's correction. *p ≤ 0.05, **p ≤ 0.01, ***p ≤ 0.001, ****p

≤ 0.0001 **(C)** Mice were surgically implanted with a cannula in the left lateral ventricle. At least 7 days later, they were sub-lethally irradiated and injected i.v. with WT T-ALL. At mid-stage disease (12-14 days later), mice were injected i.p. with DMSO (vehicle) daily for three days. On days 1 and 3 of vehicle treatment, mice received 5 microliter intraventricular infusions of isotype (47 microgram) or anti-LFA-1 and anti-VLA-4 (26 micrograms each) blocking antibodies. 24h after the last dose, tissues were isolated and T-ALL cells were enumerated. Total number of WT T-ALL cells in the CNS. Bars represent geometric mean and geometric SD. **(D)** Integrin RNA expression in blood or bone marrow biopsies from patients with T cell acute lymphoblastic leukemia, from [13]. Y-axis: Normalized expression of each integrin gene. (By comparison, average expression of *NOTCH1* is 38, average expression of *CXCR4* is 191.) X-axis: CNS grade of each patient at diagnosis. (CNS1, no detectable blasts in the cerebrospinal fluid; CNS3, at least 5 white blood cells/ml cerebrospinal fluid with detectable blasts, or other signs that leukemia has spread to the CNS.) CNS1, 183 patients; CNS2, 60 patients; CNS3, 18 patients. Statistics comparing CNS1 and CNS3 were calculated on normalized data using unpaired t-test with Welch's correction. **p ≤ 0.01 (mean *ITGB1* CNS1 27.6, CNS3 21.3).

| Integrin | WT (CRISPR) Counts | DKO (CRISPR) Counts | Ratio DKO:WT | padj |
|---|---|---|---|---|
| Itgad | 24.2 | 30.5 | 1.26 | 0.9670 |
| Itgae | 52.6 | 56.1 | 1.07 | 0.9815 |
| Itgal | 3586.0 | 2731.3 | 0.76 | 0.6881 |
| Itgam | 358.7 | 158.4 | 0.44 | 0.4571 |
| Itgav | 533.0 | 526.5 | 0.99 | 0.9956 |
| Itgax | 5.6 | 0.0 | 0.00 | 0.7309 |
| Itga1 | 0.2 | 0.9 | 4.20 | NA |
| Itga2 | 3.5 | 1.7 | 0.47 | 0.9573 |
| Itga2b | 18.3 | 27.9 | 1.52 | 0.8436 |
| Itga3 | 0.3 | 0.0 | 0.00 | NA |
| Itga4 | 1324.9 | 759.8 | 0.57 | 0.4914 |
| Itga5 | 1.9 | 0.0 | 0.00 | NA |
| Itga6 | 461.9 | 280.7 | 0.61 | 0.4029 |
| Itga7 | 8.9 | 4.9 | 0.55 | 0.9270 |
| Itga8 | 0.0 | 1.1 | Inf | NA |
| Itga9 | 4227.2 | 5269.3 | 1.25 | 0.7047 |
| Itga10 | 9.2 | 6.1 | 0.67 | 0.9540 |
| Itga11 | 1.9 | 1.1 | 0.58 | NA |
| Itgb1* | 2896.9 | 1529.6 | 0.53 | 0.0018 |
| Itgb2 | 4474.9 | 5358.8 | 1.20 | 0.7780 |
| Itgb3 | 618.5 | 684.0 | 1.11 | 0.9731 |
| Itgb4 | 12.9 | 7.2 | 0.56 | 0.8878 |
| Itgb5 | 114.0 | 85.2 | 0.75 | 0.9073 |
| Itgb6 | 0.0 | 1.9 | Inf | NA |
| Itgb7* | 7368.3 | 12030.5 | 1.63 | 0.0645 |
| Itgb8 | 0.3 | 0.0 | 0.00 | NA |

**S. Table 1. Integrin expression by CRISPR-WT and CRISPR-DKO T-ALL**

RNA-Sequencing was performed on CRISPR-WT and CRISPR-DKO T-ALL cells isolated from the CNS of mice at mid-stage disease that had received 1:1 DKO:WT T-ALL. Normalized read counts of integrin genes are shown, average of $n = 4$ biological replicates. Residual *Itgal* and *Itgb1* expression likely reflects truncated transcripts, as no surface protein remains. Statistics calculated using DESeq2 and corrected for multiple comparisons. *padj $\leq 0.1$
